# Multi-modal refinement of the human heart atlas during the first gestational trimester

**DOI:** 10.1101/2024.11.21.624698

**Authors:** Christopher De Bono, Yichi Xu, Samina Kausar, Marine Herbane, Camille Humbert, Sevda Rafatov, Chantal Missirian, Mathias Moreno, Weiyang Shi, Yorick Gitton, Alberto Lombardini, Ivo Vanzetta, Séverine Mazaud-Guittot, Alain Chédotal, Anaïs Baudot, Stéphane Zaffran, Heather C. Etchevers

## Abstract

Forty first-trimester human hearts were studied to lay groundwork for further studies of principles underlying congenital heart defects. We first sampled 49,227 cardiac nuclei from three fetuses at 8.6, 9.0, and 10.7 post-conceptional weeks (pcw) for single-nucleus RNA sequencing, enabling distinction of six classes comprising 21 cell types. Improved resolution led to identification of novel cardiomyocytes and minority autonomic and lymphatic endothelial transcriptomes, among others. After integration with 5-7 pcw heart single-cell RNAseq, we identified a human cardiomyofibroblast progenitor preceding diversification of cardiomyocyte and stromal lineages. Analysis of six Visium sections from two additional hearts was aided by deconvolution, and key spatial markers validated on sectioned and whole hearts in two- and three-dimensional space and over time. Altogether, anatomical-positional features including innervation, conduction and subdomains of the atrioventricular septum translate latent molecular identity into specialized cardiac functions. This atlas adds unprecedented spatial and temporal resolution to the characterization of human-specific aspects of early heart formation.

## Introduction

The heart is the first permanent organ to perform a vital function in all vertebrate embryos and many invertebrates (Maldonado et al., 2019). Its anatomy and output, however, vary widely across species. Understanding the normal development of human heart cells and tissues is essential for improving the diagnosis, treatment and prognosis of congenital heart defects (CHDs), the most common malformations at birth.

Due to difficulties studying human tissue directly, much of our knowledge about molecular, cellular, and morphogenetic processes driving heart development comes from animal models. While severe CHDs are often established by the sixth week of gestation, leading to spontaneous abortion, other serious or fatal defects emerge in heart regions influenced by human-specific factors like shear stress, growth, and anatomical changes that modulate cardiac function through birth (Oparil et al., 1984). These factors are difficult to reproduce in popular vertebrate models.

The late embryonic and early fetal period, at the end of the first trimester, marks a critical developmental phase of cardiac remodeling, but due to limited tissue availability it is challenging to study in humans (Hikspoors et al., 2022; O’Rahilly, 1971). Integration of new findings from additional samples is necessary for a representative atlas of prenatal cardiac cell types, functions, anatomical positions and interactions across human cellular and individual diversity (Asp et al., 2019; Cao et al., 2020; Farah et al., 2024; Hou et al., 2024; Lázár et al., 2024; Leshem et al., 2024; Xu et al., 2023).

Such integration is not trivial. Fetal heart cells produce hundreds to thousands of stage-specific transcripts and paralogous genes to those defining cell types in other parts of the body or at later life stages (D’Antonio et al., 2022). Tiny percentages of cells contact fluids or populate valves, ganglia and coordinate rhythmic contraction, requiring deep, unbiased sampling to be comprehensive. Mammalian fetal hearts display physiological properties mediated by connective tissues from various developmental lineages, about which little is known in humans (Ali et al., 2014; Deng et al., 2023; Dettman et al., 1998; Dewing et al., 2022; Gittenberger-de Groot et al., 1998; Norris et al., 2009; Vrancken Peeters et al., 1999). These considerations complicate efforts to assign cell identities on the basis of prior knowledge.

To address these challenges, we analyzed human hearts between 8 to 11 post-conceptional weeks (pcw), bridging embryonic and fetal stages. We generated *de novo* molecular and anatomically informed datasets about their composition using spatial and single-nucleus transcriptomics. Integrating published atlases, we also identified and validated novel markers, intermediates in lineage diversification, and features of the atrioventricular septation complex.

This refined atlas enhances understanding of cellular differentiation, function and spatial organization in the developing human heart.

## Results

### Increasing the resolution of biologically meaningful human cardiac cell types

To characterize even rare cells, we analyzed a total of 40 human embryonic and fetal hearts of 6.5-12.3 post-conceptional weeks (pcw) with careful anatomy, histology (Supplementary Figure S1) and complementary transcriptomics approaches, followed by spatial validation of transcript and protein expression in sliced and/or intact organs.

### Cell type assignment at single-nucleus resolution

To preserve cellular diversity among heterogeneous populations, we used single-nucleus RNA-sequencing (snRNAseq). Nuclei were isolated from whole hearts with proximal great vessels from one female (8.6 pcw) and two male (9.0 pcw and 10.7 pcw) fetuses (Supplementary Materials and Methods; Fig. 1A). We analyzed snRNAseq transcriptomes for 18,129, 8,750, and 22,348 individual nuclei respectively, and integrated the 49,227 fetal cardiac nuclei to assign group identities. Six classes comprising 21 clusters were distinguished (Fig. 1B) and their identities assigned using known markers and a post-hoc literature search for co-expressed, differentially expressed genes (DEGs) when comparing a cluster against all others (Table S1). All clusters were found in varying numbers in a hearts, such as fewer arterial smooth muscle cells (SMC) at 10.7 pcw (Fig. 1C-D). This specimen had relatively truncated distal segments of the outflow tract (Fig. S1).

**Figure 1.**
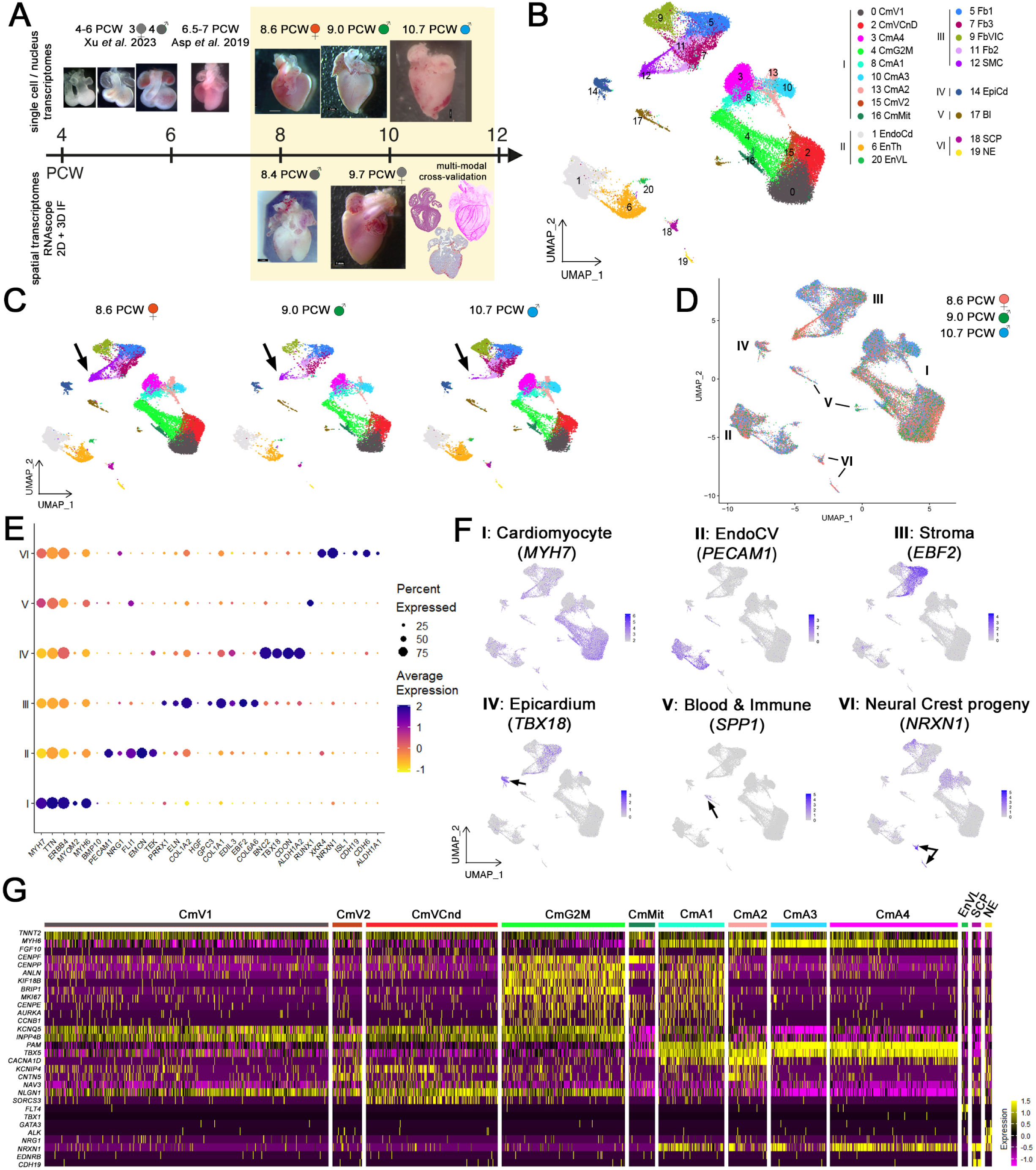
Profiling human fetal heart development at the resolution of individual nuclei (**A**) Samples of nuclei were derived from three whole dissociated hearts to generate distinct snRNAseq datasets at 8.6 post-conceptional weeks (pcw, confirmed XX genotype), 9.0 pcw (XY) and 10.7 pcw (XY). Samples of spatial transcriptomic analysis were derived from two whole snap-frozen hearts to collect cryosections and generate Visium datasets at 8.4 pcw (XY, 2 sections) and 9.7 pcw (XX, 4 sections). (**B**) Integrated UMAP representation of 49,227 profiled nuclei coloured by cell type. (**C**) UMAP representation of profiled nuclei as in (B) but separated by sample. Arrows indicate cluster 12 (SMC) in each, but that all cell types are present at each sample. (**D**) UMAP plots per sample indicating that all cell classes are also represented in each sample. (**E**) Dot plot of top marker genes for each Class (y axis: cardiomyocytes (I), endocardiovascular cells (II), stroma (III), epicardium (IV), blood (V) and neural crest progeny (VI). The size of the dot represents the percent of nuclei with transcripts at non-zero levels, and color intensity represents average log-normalized expression of the gene where relative abundance of typical markers is indicative of cell class. (**F**) UMAP feature plots of representative gene expression in Classes I to VI for *MYH7* (cardiomyocytes), *PECAM1* (endothelial and endocardial cells), *EBF2* (stroma), *TBX18* (epicardium), *SPP1* (immune cells) and *NRXN1* (neural crest). (**G**) Heatmap showing selected genes expressed in the nine cardiomyocyte clusters of Class I compared with the minority populations of lymphatic endothelium, Schwann cell precursors and neuroendocrine cells. Each column displays gene expression of an individual cell and genes are listed in the rows.

Markers such as *MYH6* and *MYH7* (Lu et al., 2022) for cardiomyocytes (I), *POSTN* and *PECAM1* for cardiovascular endothelial cells (II), and *EBF2 and PRRX1* for interstitial fibroblasts and smooth muscle for stroma (III), allowed grouping of functionally related clusters into classes. Epicardial (IV), blood and immune cell (V) and neural crest-derived (VI) cells were also designated classes (Fig. 1D-F).

### Iterative refinement of annotations

Class I cardiomyocytes (29,233 nuclei, 59.4% of total; Table S1, tab 2) broadly expressed *MYH6* and *MYH7* (Fig. 1E-G). Their relative abundance distinguished subclasses of atrial (CmA1-A4; *MYH6*^hi^) and ventricular chambers (CmV1, CmV2, CmVCnD; *MYH7*^hi^), respectively, reflecting histological (Fig. S1) and functional differences. Significant DEGs from each class and within- or between-class DEGs, are available in Table S1.

The CmG2M cluster (8.3% of class) had higher representation of G2M transition characteristics, including significant DEGs encoding the centriolar assembly proteins *CENPF* and *CENPP*; *ANLN* (cytokinesis); *KIF18B* and *KIF15* (kinesin superfamily); and *BRIP1*, a BRCA1-interacting helicase. CmMit represented mitotic cardiomyocytes (3%). Its significant DEGs encoded *MKI67*, centriolar *CENPE*, the cell cycle kinase *AURKA*, and the mitosis-initiating *CCNB1*. *MYH7* and *TTN* were among DEGs of the ventricular cluster CmV1 (32.2%, Fig. 1G) but CmV1 also uniquely co-transcribed *KCNQ5* (potassium channel Kv7.5) and *INPP4B* (a PI3K signaling suppressor; Fig. S2A). This combination may underscore the specific electrical properties of ventricular cardiomyocytes.

CmA2 and CmV2 spanned atrial and ventricular subclasses (4.4 and 3.4% of Class I cells; Fig. 1D, Table S1). With *PAM*, *TBX5* and *MYH6* as DEGs, CmA2 also expressed the highest atrial levels of *CACNA1D* (Table S1, tab 7). *CACNA1D* encodes the L-type voltage-activated calcium channel CaV1.3 of the sinoatrial node (van Eif et al., 2019). CmV2, like CmVCnD, expressed *KCNIP4* (a voltage-gated potassium channel interacting protein) but like CmA2, expressed the AV bundle adhesion contactin *CNTN5* (Fig. 1G) (Kanemaru et al., 2023; Smirnov et al., 2018). CmVCnD differed from CmV2 and other ventricular cardiomyocytes (Table S1, tab 6) by *NAV3* (neuron navigator 3), an axon guidance cue required for zebrafish heart morphogenesis (Lv et al., 2022). Other CmVCnD markers included *NLGN1*, encoding a ligand for post-synaptic neurexins of neuromuscular junctions (Ramesh et al., 2021) and *SORCS3*, a neurotrophin uptake receptor necessary for post-synaptic vesicular trafficking (Oetjen et al., 2014) (Fig. 1G). CmA2, CmV2 and CmVCnd distinguished cell states and/or locations of the developing human pacemaker (sinoatrial and atrioventricular nodes) and conduction system.

Rare clusters were assigned identities by comparing DEGs first to all other heart nuclei, then to the other clusters within their class, and leveraging Enrichr-KG (Evangelista et al., 2023) for significantly enriched annotations from the Descartes human fetal atlas (Cao et al., 2020), the Human Gene Atlas (Su et al., 2004), the Tabula Sapiens (The Tabula Sapiens Consortium, 2022), the KEGG 2021 Human, and the Gene Ontology Biological Process databases (Fig. S2B). For example, neural crest-derived Class VI (*NRXN1*, *CDH6*) subdivided into neuroendocrine (NE; 227 nuclei; Fig. 1E-G, Fig. S2B) and Schwann cell precursor (SCP; 309 nuclei; Fig. 1G, Fig. S2C) clusters. *NRG1*, *GATA3* and *ALK* (Boeva et al., 2017) distinguished NE from SCP (*EDNRB*, *CDH19*) (George et al., 2018).

The smallest EnVL (venous-lymphatic endothelium) Class II cluster of 210 cells (Fig. 1B, G) co-expressed established markers like *FLT4* (*VEGFR3*) and *TBX1* (Fig. 1G; Table S1) (Wang et al., 2023). Although intentionally rinsed away during collection, blood-derived Class V (655 nuclei) included fetal hemoglobin-expressing (*HBG1*/*HBG2*) erythrocytes, *RUNX1*-expressing immature and *PTPRC* (CD45)-expressing differentiated hematopoietic progeny, and *SPP1*-positive macrophages (Fig. 1D-E).

### Integration of first-trimester cardiac atlases at cellular resolution identifies an early cardiomyofibroblast progenitor

We integrated our snRNAseq data with two human heart atlases derived from single-cell (sc)RNAseq of whole human embryos aged 4-6 pcw (8,266 cells) (Xu et al., 2023) and a 6.5-7 pcw heart (3,717 cells) (Asp et al., 2019) to analyze transcriptomic changes over time (Fig. 2A). Similar integration of scRNAseq and snRNAseq datasets has enabled identification of minority progenitors in mouse skeletal muscle (McKellar et al., 2021).

**Figure 2.**
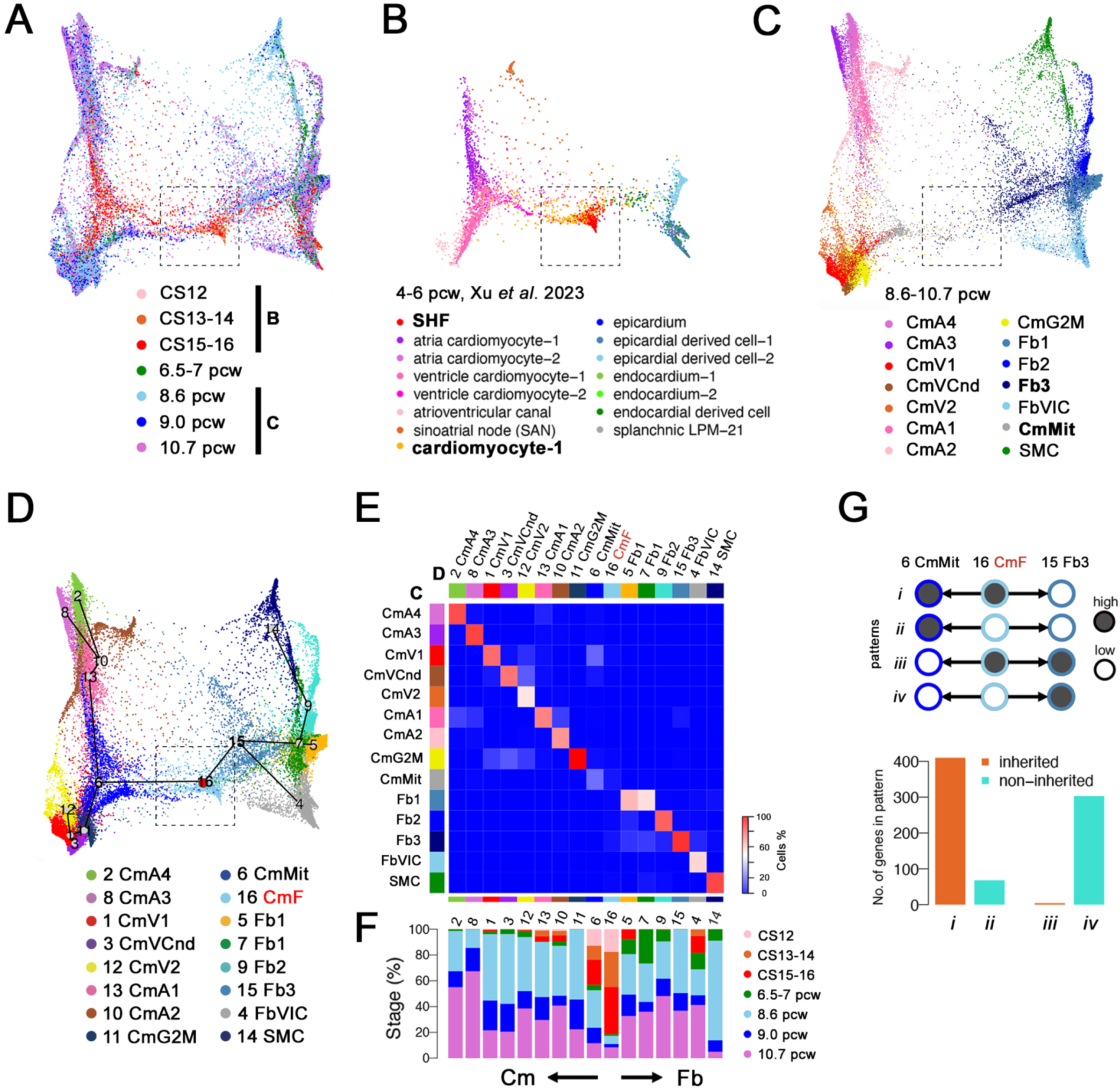
Trajectory analysis of Classes I and III over time suggests a transient cardiomyofibroblast progenitor present at 4-6 pcw (**A**) PHATE visualization of cardiomyocyte and stromal cells, colored by developmental stage, from cardiac single-cell RNA-seq data of seven human embryos at Carnegie stage (CS)12, CS13-14 and CS15-16 (Xu et al., 2023), one heart at 6.5-7 pcw (Asp et al., 2019), and this study’s snRNAseq data from hearts at 8.6, 9.0 and 10.7 pcw. Boxed area in **A-D** delimits a cell population with greatest contribution from early stages. (**B**) Subset of PHATE plot from A, showing scRNA-seq data at 4-6 pcw, colored by original clusters from Xu et al., 2023. (**C**) Subset of PHATE plot from **A** showing snRNAseq data at 8.6, 9.0 and 10.7 pcw, colored by integrated clusters (cf. Fig. 1B). (**D**) Superimposition of Slingshot trajectory on PHATE plot in **A-C**, predicting cluster lineage bifurcations (black lines) from cluster 16, the “cardiomyofibroblast” (CmF; dashed square **A-D**). (**E**) Confusion matrix of percentages of cell types from our snRNAseq clusters (**C** on y axis) against new clusters after dataset integration (**D** on x axis). Fewer than 20% of the cells in cluster 16 (CmF) persist after 8.6 pcw. (**F**) Histogram showing percentages of cells/nuclei from each stage in each cluster from **D**, aligned with **E**’s x-axis. Fb2, SMC and CmVCnd cell types appear after 6.5 pcw, while CmA3, CmG2M and Fb3 were newly identified after 8.6 pcw. (**G**) Top panel shows four inheritance patterns (1-4) of the trajectory from cluster 16 (CmF) to 6 (CmMit) and 15 (Fb3). Bottom panel: DEGs of A>B defined as A > 0.5 and A/B > two-fold. 1: CmF and CmMit > Fb3 (inherited DEGs). 2: CmMit > CmF and Fb3 (non-inherited). CmF and Fb3 > CmMit (inherited DEGs). Fb3 > CmF and CmMit (non-inherited). Patterns 1 and 4 best fit the data, suggesting that CmF resembles a cardiomyocyte more than a fibroblast.

Cardiomyocytes and stromal cells in the mouse derive from common mesodermal progenitors that begin lineage restriction as early as gastrulation (Lescroart et al., 2014). To search for evidence of similar human fate engagement, we integrated cells assigned to stromal or cardiomyocyte classes (I and III, Fig. 1D) between 5 pcw (Carnegie Stage [CS]13) and 10.7 pcw. We then performed dimensional reduction with Potential of Heat-diffusion for Affinity-based Trajectory Embedding (PHATE), a diffusion-based manifold learning method (Moon et al., 2019) (Fig. 2A-C; Fig. S3C-D). The new clusters corresponded on a one-to-one or one-to-many basis with our snRNAseq cell type assignments (Fig. S3A-B). Some assignments were nearly exclusive to later cells, including clusters 8 (CmA3) and 2 (CmA4) as well as a stromal progenitor in cluster 15 (Fb3) (Fig. 2D-F). In contrast, cells from the 6.5-7 pcw heart (Asp et al., 2019) were relatively impoverished in cardiomyocytes and their precursors (Fig. 2F), perhaps due to technical loss of larger cells *versus* size-agnostic nuclei.

Not all clusters were equally populated by stage; one in particular appeared more represented at CS15-16 (6 pcw) than at later stages (Fig. 2A-C, box). Slingshot trajectory analysis (Street et al., 2018) superimposed on this new projection traced classes I and III back to a common cardiomyofibroblast (cluster 16, CmF), mostly depleted after 6 pcw (Fig. 2D). A confusion matrix between the full scRNAseq/snRNAseq dataset (Fig. 2E) and our original clusters (Fig. 1) showed little overlap between CmF and later clusters belong to cardiomyocyte or stromal classes, although some CmF persisted even at 10.7 pcw (Fig. 2C, F).

CmMit (cluster 6) and Fb3 (cluster 15) were the nearest class-specific progenitors to CmF (Fig. 2D). To identify which more resembled parental CmF, we classified all significant DEGs into “high” or “low” bins, scoring how these categories were inherited over the trajectory from cluster 16 to 6 or to 15 during differentiation. Most CmF genes (Table S2) were maintained from CmF to CmMit but newly transcribed in Fb3, suggesting closer lineage ties to proliferating cardiomyocyte progenitors (Fig. 2G).

In summary, our snRNAseq data segregated into six classes and 21 cell clusters, highlighting relevant cardiomyocyte identities, rare cell types, anatomical and developmental features. Integrating existing data, we identified new progenitor states and modeled lineage relationships. These findings enhance current understanding of human cardiac cell diversification and organization.

### Spatial transcriptomics of multiple 8.4 and 9.7 pcw heart sections

To add positional context, we undertook 10x Genomics Visium spatial transcriptomics (ST) analysis on two sections from an 8.4 pcw male heart and four sections from a 9.7 pcw female heart (Fig. 1A), assessing over 12,000 regularly spaced tissue spots (Supplementary Materials and Methods). We identified eight clusters in 8.4 pcw sections and 6-8 at 9.7 pcw (Figs. S4-9). Cluster information was then projected onto each section at appropriate coordinates. Histological landmarks (Fig. S1) and known markers permitted us to assign cluster identities: for example, compact or trabecular *MYH7*+ ventricular cardiomyocytes (*HEY2 and IRX3*, respectively). *MYH6*+ atrial cardiomyocytes expressed *BMP10* and *TBX5* on the right and *PITX2* on the left. Arterial smooth muscle enriched in *ELN* and *MYH11* was identified at both stages, as were discrete clusters of valvular (*SCX*, *PENK*) or subvalvular (*POSTN, PRRX1*) stroma. Full lists of markers in individual section clusters are provided (Table S3). Integration of these datasets distinguished 14 clusters (abbreviated SIC to distinguish from snRNAseq clusters), improving cell type comparisons across time and space (Fig. 3A, B, and Fig. S10; Table S4). Left and right, compact and trabecular cardiomyocyte clusters expressed expected genes (Fig. 3B, C), but integration enabled better discrimination of non-contractile cell types with spatial coherence (Table S4).

**Figure 3.**
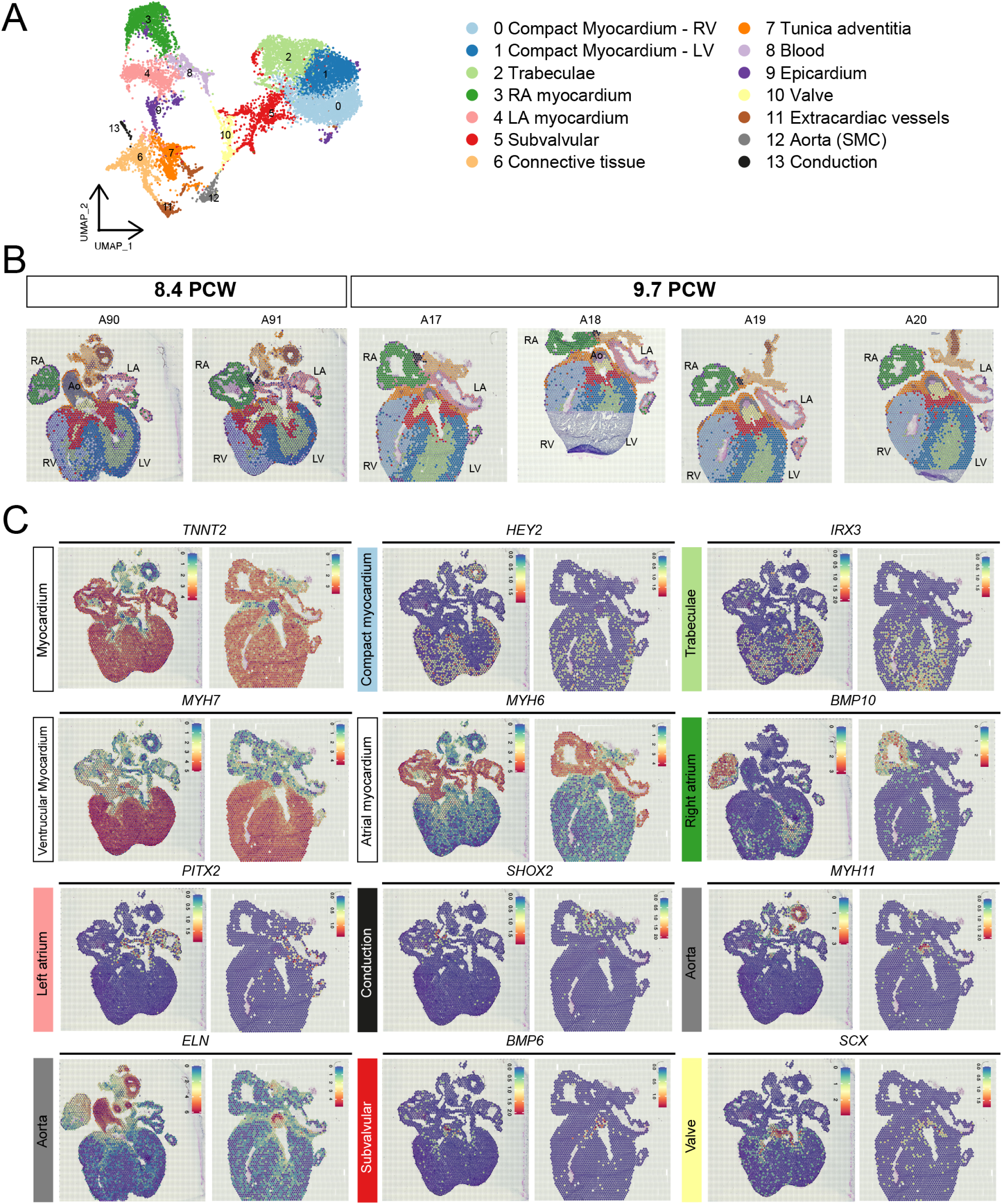
Spatial transcriptomics (ST) in 8.4 pcw and 9.7 pcw human heart sections show regionalized distributions of cardiac cell types (**A**) Seurat UMAP plot of integrated spatial transcriptomic data from two 8.4 postconceptional week (pcw) and four 9.7 pcw heart sections. Each dot represents a tissue-covered spot from the Visium spatial capture slides. Fourteen clusters corresponding to cardiac cell types are listed. (**B**) Visualization of clustering on each heart section after ST data integration, with cluster annotations and colors matching the UMAP in A. (**C**) Spatial plots of gene expression, with highest in red and lowest in blue. RV, right ventricle; LV, left ventricle; RA, right atrium; LA, left atrium; Ao, aorta; SMC, (vascular) smooth muscle cells.

The ascending aortic SMC (SIC12; *ELN*+) was clearly distinct at both stages from thick-walled, extracardiac arteries. *Tunica adventitia* (SIC7; *COL1A1*+) of the ascending aorta and coronary arteries was further abluminal. SIC6 included both myofibroblasts (*PRRX1*, *DCN, FN1, COL1A2, COL3A1*) and encased autonomic neurons (*NEFL*, *TUBB3*, *PRPH*, *PHOX2B*). The valves (SIC10) expressed *SCX* (Fig. 3C) and *POSTN*, but SIC5 and SIC12 were also strongly enriched in *POSTN*. Epicardial identity (SIC9) was marked by *TBX18* and *ITLN1* (Sacks and Fain, 2007). The cardiac conduction system, with the fewest spots, corresponds in these sections to the sinoatrial node (SIC13; *SHOX2*, *TBX3*), demonstrating improved cell type annotation by adding ST (Table S4; Fig. 3A-C).

Immunofluorescence validated MYL7, MYH6 and MYH7 protein distribution (Fig. 4). Myosin is a hexamer composed of four light and two heavy chains, necessary for contractility. Myosin regulatory light chain 7 (encoded by *MYL7*) is a human atrial-specific gene, expressed from 8 pcw until after birth (Hailstones et al., 1992). While *MYL7* was initially pan-cardiac (8.4 pcw, Fig. 4A-B), it was predominantly atrial by 9.7 pcw (Fig. 4C), consistent with its immunoreactivity at 9.0 pcw (Fig. 4D) and confirming *in situ* hybridization at the same stage (Hailstones et al., 1992). MYH6 and MYH7 were detected in the atria and the ventricles at 9.0 and 12.0 pcw respectively (Fig. 4E-L). As predicted from avian data (Teichmann and Kessel, 2004), *BMP10* was enriched in ventricular trabeculae and the right atrium (Fig. 4M), as confirmed by RNAscope at 9.0 pcw (Fig. 4N-P).

**Figure 4.**
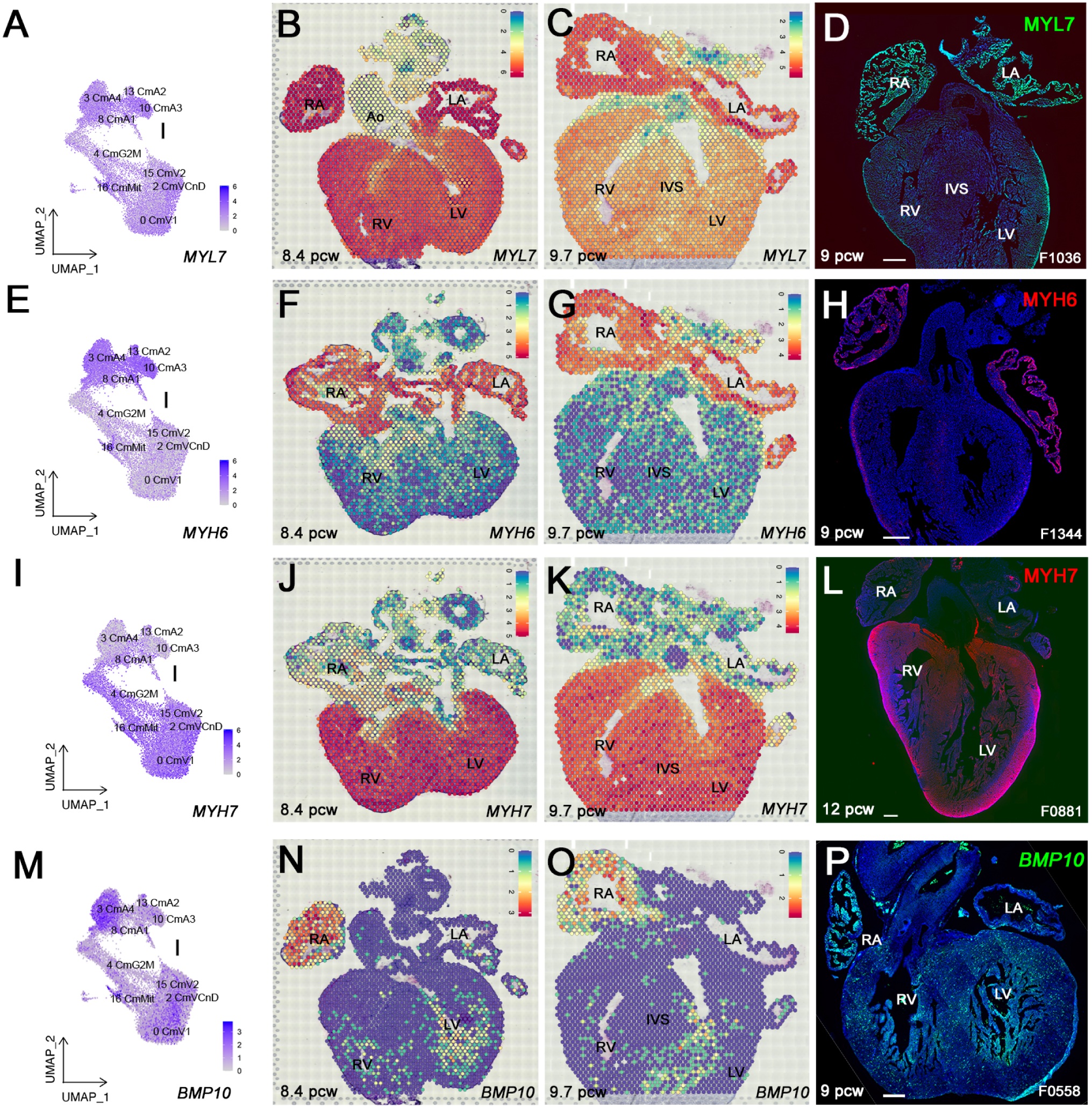
Corroboration of Class I marker expression (First column: **A, E, I, M**) UMAPs showing gene expression of *MYL7*, *MYH6*, *MYH7* and *BMP10* in cardiomyocyte Class I, indicating atrial and ventricular subclasses. (**B, C**) Spatial transcriptomic (ST) plots of *MYL7* transcript distribution. (**D**) Immunofluorescence of MYL7, with greater atrial than ventricular expression. (**F, G**) ST plots of *MYH6*. (**H**) MYH6 more strongly immunostained atria than ventricles. (**J, K**) ST plot distribution of *MYH7*. (**L**) Immunofluorescence of MYH7 in ventricular myocardium at 12.0 pcw. (**N, O**) ST plots of *BMP10* . (**P**) *BMP10* transcripts detected by RNAscope in ventricular trabeculae and pectinate muscles of the right atrium . (**B, F, G, J, K, N, O**) Highest expression in red and lowest in blue. Ao, aorta; IVS, interventricular septum; LA, left atrium; LV, left ventricle; RA, right atrium; RV, right ventricle. Scale bars: 250µm.

Integrated spatial transcriptomics refined our fetal heart atlas by adding positional context, enabling finer identification of certain cell types and expression dynamics over time.

### Deconvolution of spot transcriptomes

Each spot in sequencing-based ST may contain distinct cell types in varying proportions. To measure this heterogeneity, we deconvoluted each spot using the Robust Cell Type Decomposition (RCTD) method (Cable et al., 2022), referencing our integrated snRNAseq data. Relative cell-type proportions were projected back onto the six heart sections, turning each spot into pie charts of relative snRNAseq cluster proportions (Fig. 5, Fig. S11-16).

**Figure 5:**
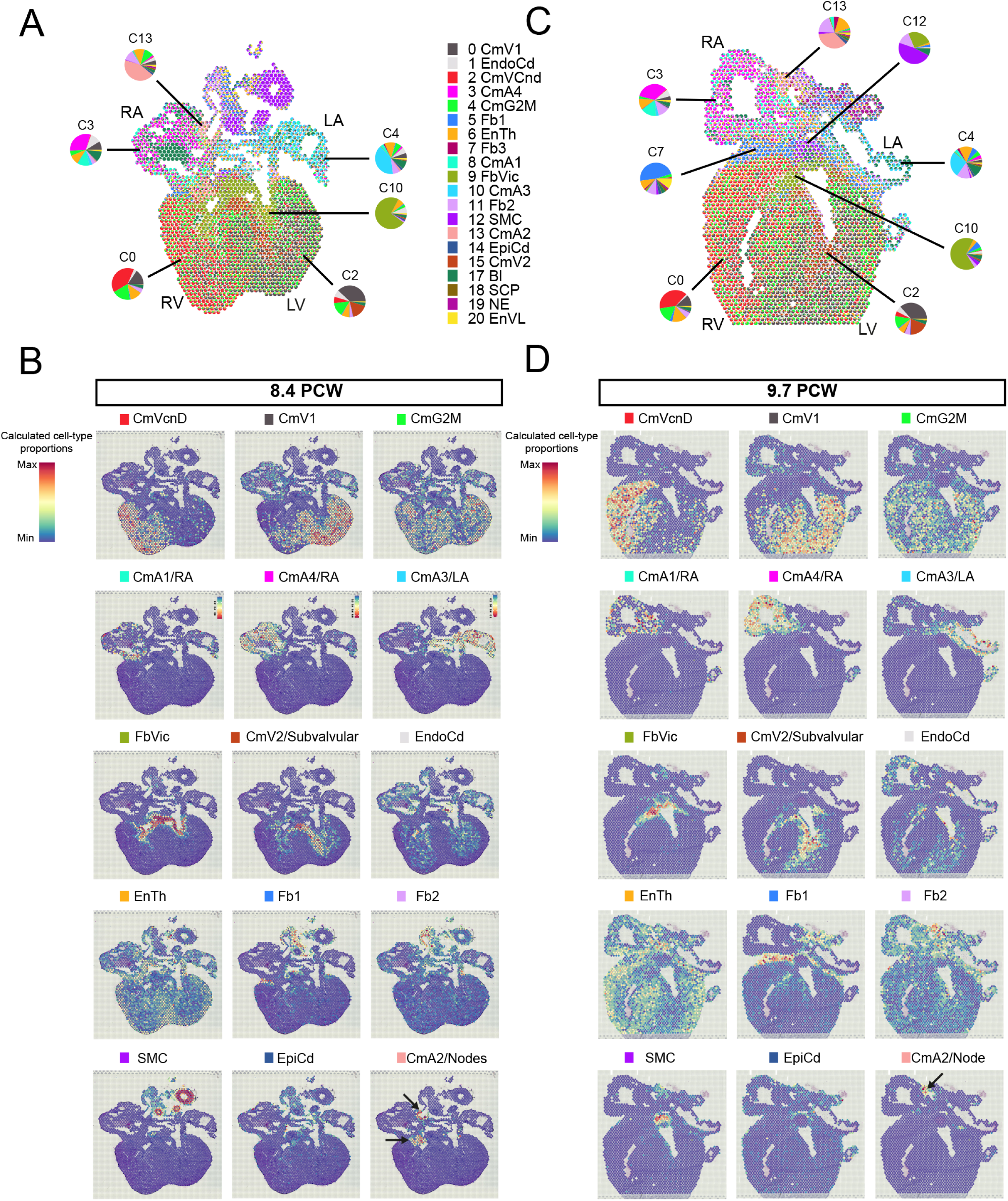
Deconvolution of heart ST data using snRNAseq reference reveals transcriptional heterogeneity in ST spots (**A, C**) Deconvolution of spots of 8.4 pcw heart section A91 (**A**) and 9.7 pcw section A17 (**C**) using RCTD. The proportion of each cell type (cluster) from integrated snRNAseq data (Figure 1) is represented in pie charts for each spatial spot. Large pie charts represent the average proportion of each cell type in spatial clusters C0, C2, C3, C4, C10 and C13 from Figure 3B. (**B, D**) Cell-type proportions for CmVCnD, CmV1, CmG2M, CmA1, CmA4, CmA3, FbVIC, CmV2/subvalvular, EnTh, EndoCd, Fb1, Fb2, SMC, EpiCd and CmA2/node in spots of 8.4 pcw section A91 (**B**) and 9.7 pcw section A17 (**D**). Highest proportion is in red, lowest in blue. Note the consistency between snRNAseq cell types and their predicted spatial localization.

Figure 5 highlights 8.4 pcw heart section A91 and 9.7 pcw heart section A17. Pie charts depict snRNAseq cell types per spot (Fig. 5 A,C). Deconvolution results were further averaged in each of the 14 ST clusters across the 6 integrated ST datasets (Fig. 5 A,C; Table S5) as large pie charts with mean snRNAseq cell type proportions in SIC0 (right ventricular compact myocardium), SIC2 (trabeculae), SIC3 (right atrial cardiomyocytes), SIC4 (left atrial cardiomyocytes), SIC7 (*tunica adventitia*), SIC10 (valves), SIC12 (aorta, vascular SMC) and SIC13 (conduction system). For example, at 8.4 pcw (Fig. 5A), SIC0 comprised on average 40% CmVCnD, 15% CmV1, 19% CmG2M and 12% EnTh profiles among others, while SIC10 contained on average 73% FbVIC cells, 6% EndoCd and 7% of EnTh cells.

Deconvolution results for sections A91 (8.4 pcw) and A17 (9.7 pcw) are shown in Fig. 5B D, with estimated relative proportions of the following clusters from integrated snRNAseq data: CmVCnD, CmV1, CmG2M, CmA1, CmA2, CmA3, CmA4, CmMit, EnTh, EndoCd, Fb1, Fb2, FbVIC, SMC and EpiCd. Similar proportions were observed across all sections (Figs. S11-16; Table S5).

### Discrete chamber identities by the third gestational month

Deconvolution refined integrated snRNAseq cluster identities. For example, at 8.4 pcw (A91), right atrium spots (SIC3) averaged 30% CmA4 and 12% CmA1, while left atrium spots (SIC4) averaged 40% CmA3 but also 5% CmV1, 6% CmG2M and 3.5% CmA1 (Table S5). CmA4 correlated to right atrial identity and CmA3 with left, but their purported progenitor CmA1 (Fig. 2D) appeared in both atria (Fig. 5B, D).

The CmA2 cluster mapped to sinoatrial and atrioventricular nodes, which for the former was observed across multiple sections at each stage (Fig. 5B, arrows; Figs. S11-16). CmVCnD cluster dominated in the right ventricle, while CmV1 was more prominent in the left ventricle; these identities were observed in compact and trabecular myocardium (Fig. 5B, D). CmV2 mapped to trabeculae and a septal subvalvular domain, while EndoCd designated trabecular endocardium. SMC of aorta and extracardiac arteries surrounded EnTh while Fb1 localized to the *tunica adventitia*, capping the ventricles under the atria (Fig. 5 and Figs. S11-16).

Combining snRNAseq and ST data revealed extensive cell-type heterogeneity and refined identities of snRNAseq clusters. We appended positional terms (*eg*. left/right atria, subvalvular, atrioventricular node) to snRNAseq cluster names (Fig. 5) and confirmed distinct identities of endocardium and vascular endothelia.

### Multi-omics identifies small cell clusters and rare transcripts

Integrating the two modalities allowed us to highlight cell markers not previously implicated in human cardiac development. Fibroblast and mural cell types in the mouse are functionally diverse and site-specific (Muhl et al., 2020). To identify similar specializations, we compared the combined list of genes significantly enriched in the clusters of snRNAseq stromal Class III (Table S1) with those significantly enriched in six spatial clusters corresponding to valves, subvalvular and connective tissue, *tunica adventitia*, extracardiac vessels and aortic SMC (Table S4). *PENK*, enriched in FbVIC in snRNAseq analysis, was expressed both in and between valves in ST (Fig. 6A, Fig. S17). Other valve DEGs included *BOC*, *INHBA*, *LTBP2*, *MEOX1*, *PTN*, *PTPRZ1*, *PRRX1*, *ROBO1,* and *RSPO3* (Fig. 6A, Fig. S17).

**Figure 6:**
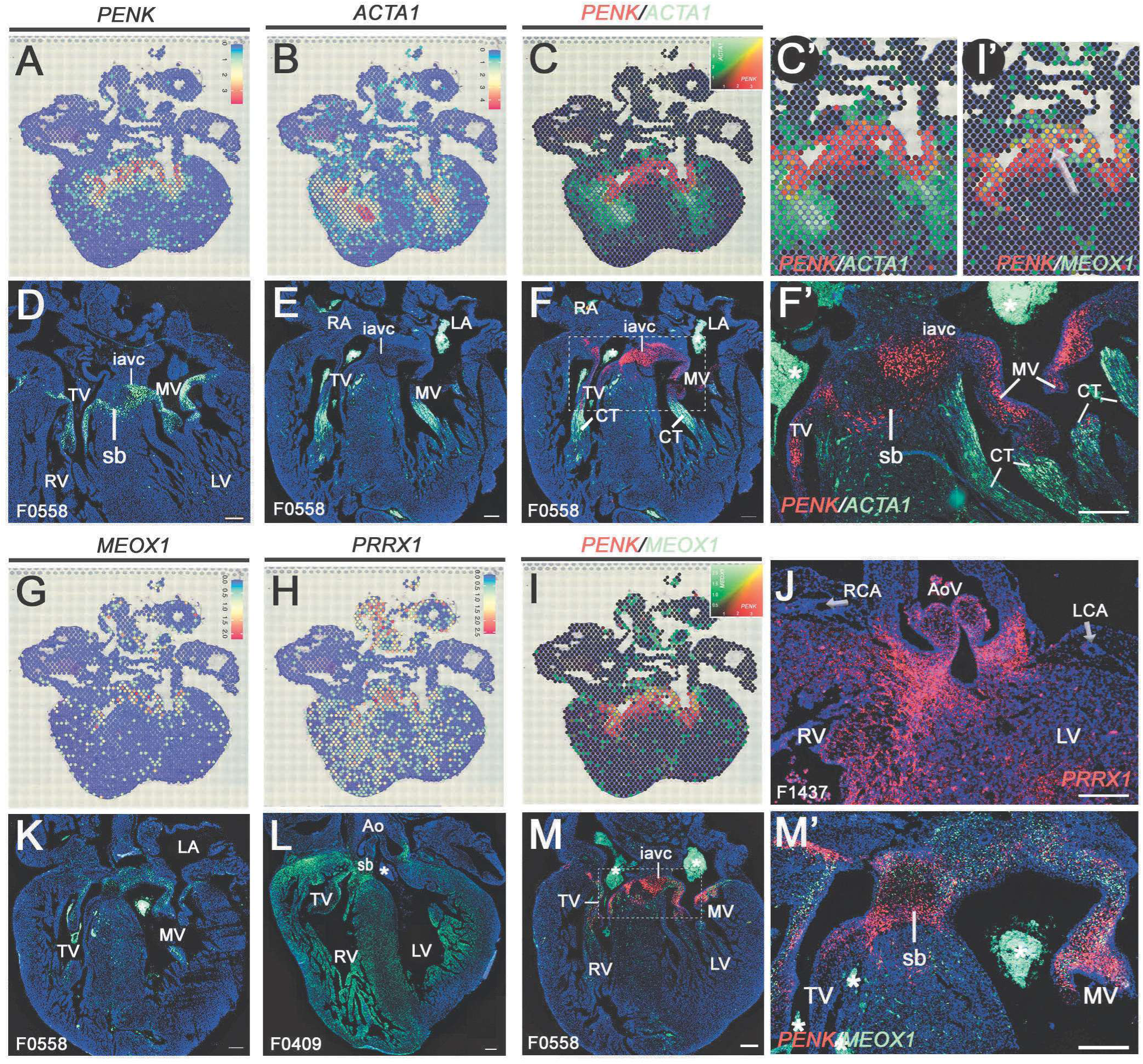
Spatial gene expression in outflow and atrioventricular valves (**A-C, G-I**) Spatial plots of stromal class transcripts in representative sections of 8.4 pcw heart, with highest expression in red and lowest in blue. (**C, I**) Co-expression plots of *PENK* (red) and *ACTA1* or *MEOX1* (green), with highest expression of each in the most saturated color and equal co-expression in yellow. (**C’, I’**) Enlargement of the atrioventricular cushion region shown in panels **C** and **I**. (**D, E**) RNAscope *in situ* hybridization of *PENK* and *ACTA1* in a 9.0 pcw heart section. (**F, F’**) Double-staining showing expression of *PENK* (red) and *ACTA1* (green) mRNAs in the same section. *PENK* is expressed in atrioventricular valve leaflets, while *ACTA1* labels chordae tendineae. (**F’**) Enlargement of the area marked by a dotted square in the adjacent section from panel F. (**J**) *PRRX1* in the aortic valve of a 7.0 pcw heart. (**K, L**) Spatial expression of *MEOX1* and *PRRX1* detected by RNAscope in a 9.0 pcw heart. (**M, M’**) Double-staining of *PENK* (red) and *MEOX1* (green) mRNAs in the same section. (**M’**) Enlargement of the area marked by a dotted square in the adjacent section from panel M. *PENK* is expressed in the inferior atrioventricular endocardial cushion, partly overlapping (arrows in **I’**) with *MEOX1*. Ao, aorta; CT, chordae tendineae; iavc, inferior atrioventricular endocardial cushion (Anderson et al., 2003); LA, left atrium; LCA, left coronary artery; LV, left ventricle; MV, mitral valve; RA, right atrium; RCA, right coronary artery; RV, right ventricle; TV, tricuspid valve; sb, subvalvular domain (septal bridge); *, blood clots. Arrows (**J)**, coronary arteries. Scale bars: 250 µm except D, 500 µm.

RNAscope analysis in additional specimens confirmed *PENK* distribution (Fig. 6D, F, F’, M, N). *ACTA1* (Actin Alpha 1, Skeletal Muscle), was the top DEG of the complementary subvalvular ST cluster (Table S4; Fig. 7). RNAscope validated *ACTA1* expression at the ventricular attachment points of the valvular leaflets and the chordae tendineae (the “heart strings”; Fig. 6E, F, F’). ST analysis predicted mutually exclusive domains for *ACTA1* and *PENK* (Fig. 6G), which we confirmed by RNAscope (Fig. 6G’). Comparison of subvalvular ST DEG to other stromal class clusters (6, 7, 10, 11, 12) found common differential expression of only two genes, both cell cycle-associated: *ANLN* and *CDK1* (Table S4). Consistently, CmG2M contributed most to the subvalvular domain, at over 19%, but CmV2, CmVCnd and CmV1 were close behind (Table S5).

**Figure 7:**
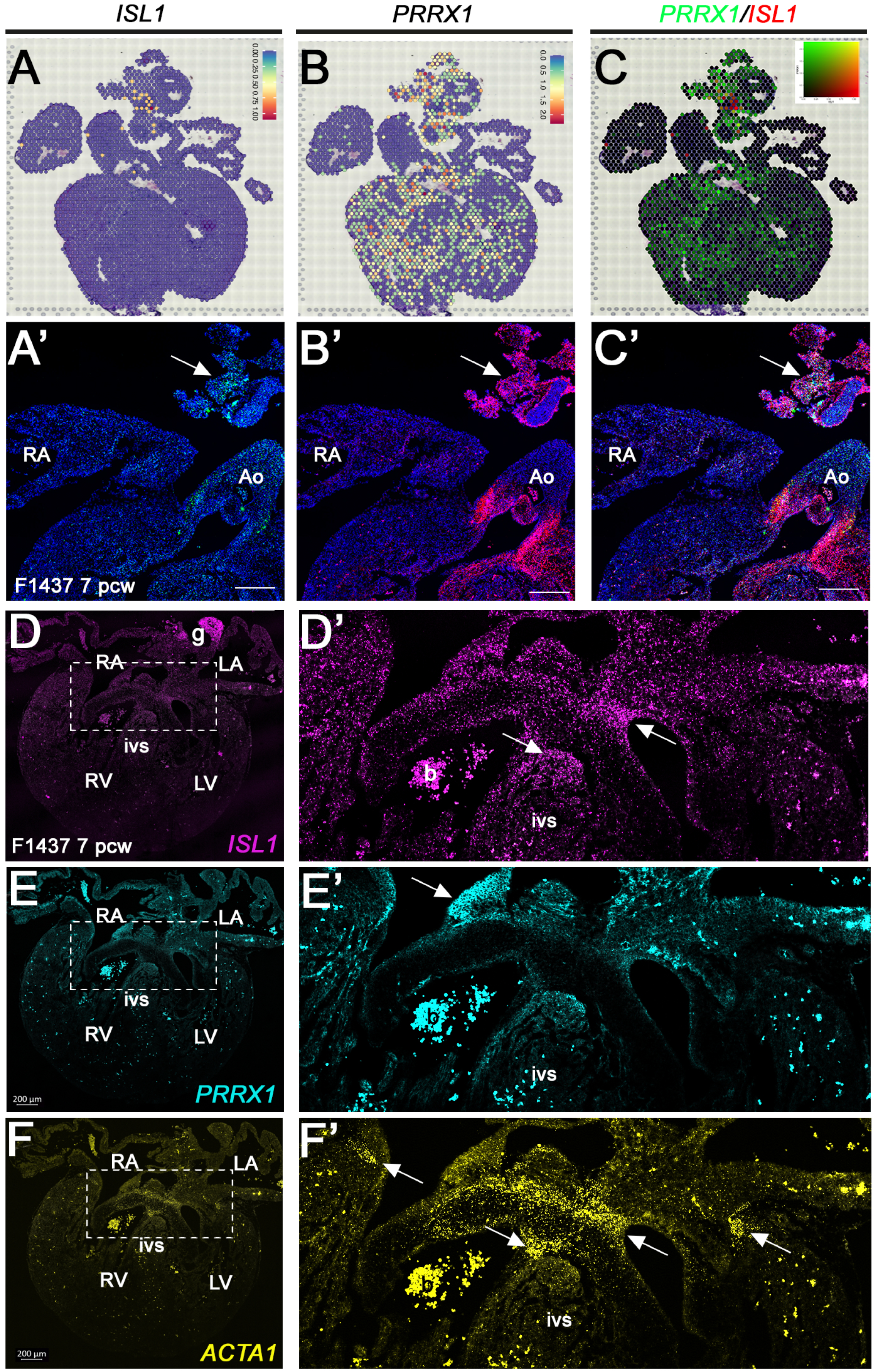
Parasympathetic ganglia and nerve markers (**A, B**) Spatial plots of transcripts enriched in conduction and nervous system on an 8.4 pcw heart section, with highest expression in red and lowest in blue. (**C**) Co-expression plots of *ISL1* (red) and *PRRX1* (green), with highest expression of each in the most saturated color and equal co-expression in yellow. **(A’, B’**) RNAscope *in situ* hybridization detected *ISL1* and *PRRX1* in the same sections. (**C’**) Complementary expression of *ISL1* (green) and *PRRX1* (red) in a parasympathetic ganglion. (D-F) *ISL1*, *PRRX1* and *ACTA1* respective RNAscope probes co-hybridized in another section from same 7 pcw heart. At atrioventricular septum intersection in box, enlarged in D’-F’, localized expression of each transcript is indicated with arrows. Ao, aorta; b, blood; ivs, interventricular septum; LA, left atrium; LV, left ventricle; RA, right atrium; RV, right ventricle. Scale bars (**A-C**): 250µm; (**D-F**): 200 µm.

Mesenchymal homeobox-1, *MEOX1*, which regulates fibrosis of injured postnatal hearts (Alexanian et al., 2021), was significantly enriched in FbVIC and Fb1 fibroblasts as shown by snRNAseq (Table S1) and transcribed in atrioventricular (AV) valves (Fig. 6G, K, M’) . *PRRX1*, a widely expressed master transcription factor associated with conduction anomalies and a top DEG characteristic of CmVCnd (Guo et al., 2021; Ke et al., 2022; Lee et al., 2022), was co-expressed with *MEOX1* in the tricuspid valve (Fig. 6I, M, M’). We confirmed that *PRRX1* was transcribed not only in AV and aortic valves, but also in the subvalvular domain (SIC5) and ventricular trabeculae (SIC2; Fig. 6H, J, L).

The connective tissue cluster SIC6 featured co-expression of many genes associated with autonomic nerve development, including *ISL1*, *TH*, *PRPH*, *PHOX2A*, *PHOX2B*, *PRRX1*, *DLK1*, *PENK*, *TBX2, GAP43*, *L1CAM*, *STMN2* and *TUBB3* (Fig. S18). RNAscope and immunofluorescence analysis confirmed complementary expression of PRPH protein and *ISL1* and *PRRX1* transcripts (Fig. 7A-C). SIC6 thus also comprises embedded cardiac ganglia. Overall, these results validated the different modes of transcriptional analysis, including deconvolution of ST spot heterogeneity with a snRNAseq reference.

### Toward a four-dimensional atlas

Cell types along reticulated blood vessels, connective tissues and nerves are represented in all organs, but can be difficult to detect in ST because of low absolute cell numbers or fine layers with distinct functions. The 536 (1.1% of total) cells assigned to neural crest-derived Class VI were not well resolved by ST outside a ganglion fortuitously sampled in section A90. The 7,224 transcriptomes of endocardial, vascular or lymphatic endothelial cells were not as well mapped as abluminal SMC and connective tissue. Although we had identified clusters corresponding to neuronal (*PRPH*+) and endothelial (*PECAM1+*) cells (Fig. 1 and Fig. 8A, D), the logic of their anatomic distributions was invisible in ST (Fig. 8B, E). Two physical dimensions are insufficient to represent the architecture of blood vessels and peripheral nerves within and surrounding the heart (Fig. 8C, F).

**Figure 8.**
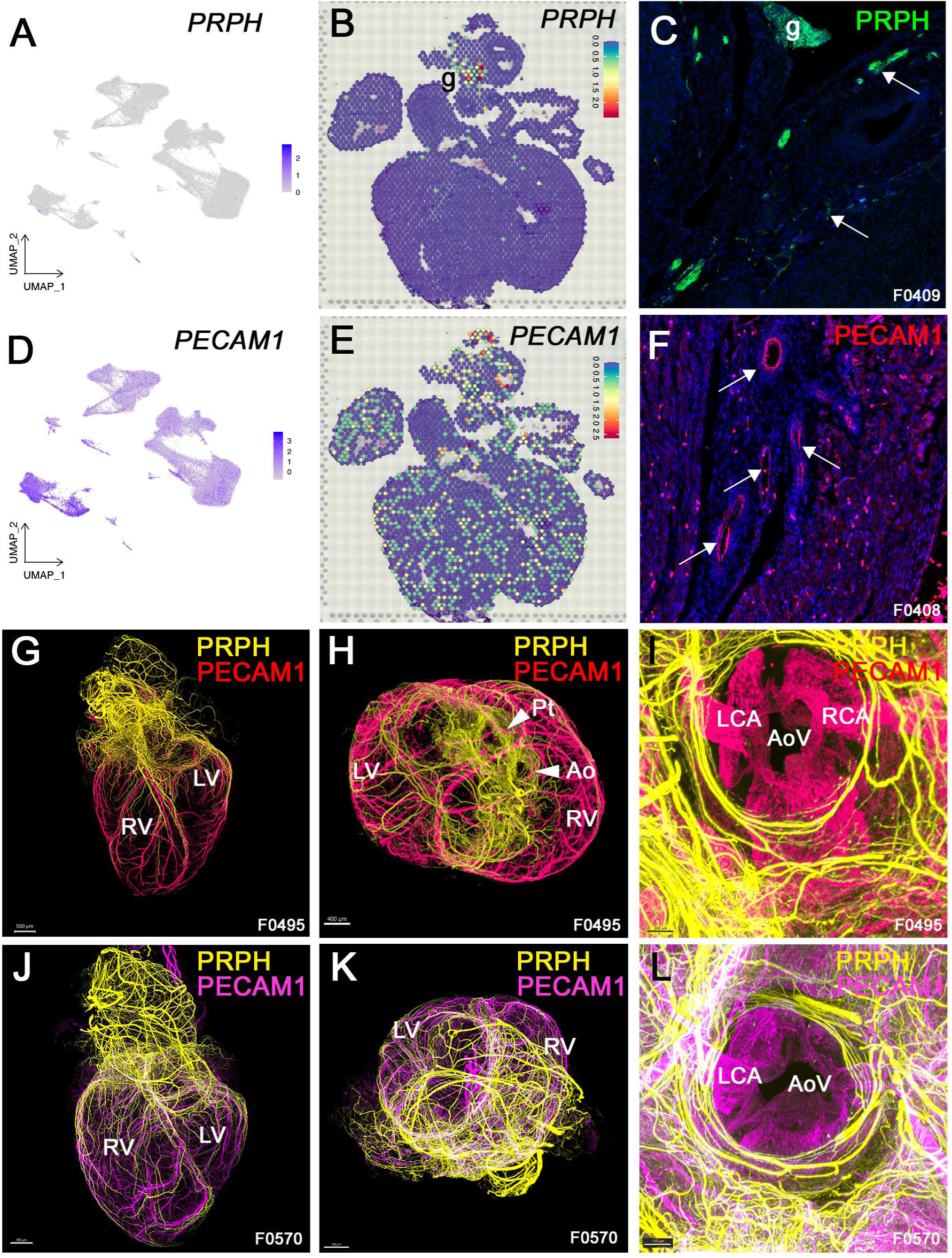
Three-dimensional reconstruction of vascular and nerve scaffolds over time (**A**) UMAP visualization of *PRPH* by snRNAseq. (**B**) Spatial plots showing *PRPH* expression in cardiac ganglion (g) on an 8.4 pcw heart section. (**C**) Immunofluorescence detection of PRPH in a 9.0 pcw heart section. PRPH is expressed in the nerves (arrows) and ganglia (g). (**D**) UMAP visualization of *PECAM1* by snRNAseq. (**E**) Spatial plots showing *PECAM1* transcripts in an 8.4 pcw heart section, with highest expression in red and lowest expression in blue. (**F**) PECAM1 (CD31) immunofluorescence in vascular endothelial cells (arrows) in a 9.0 pcw heart section. (**G-L**) Whole-mount immunofluorescence of both PECAM-1 (CD31, red [**G**] or magenta [**J**]) and PRPH (yellow) in 9.0 and 10.0 pcw hearts. (**G, L**) 3D reconstructions from light-sheet fluorescent microscopy (LSFM) image stacks show parallel networks of vessels (PECAM1+) and nerves (PRPH+) in 9.1 and 10.1 pcw hearts. (**H, K**) Top views of hearts in **G** and **J**. (**I, L**) LSFM reconstructions of the pulmonary valve showing that valve leaflets, unlike the great vessels, are not innervated. (**B, E**) Highest expression in red, lowest in blue. Ao, aorta; AoV, aortic valve; LCA, left coronary artery; LV, left ventricle; Pt, pulmonary trunk; RCA, right coronary artery; RV, right ventricle. Scale bars: (**G, J**) 500 µm; (**H**) 400 µm; (**I**) 300 µm; (**K**, **L**) 150 µm.

Whole-organ immunofluorescence after tissue clearing has enabled better sampling of peripheral nerve ramifications and lacrimal gland development over time (Belle et al., 2017; Blain et al., 2023). We used whole-organ immunofluorescence to examine spatial arrangements of blood vessels and peripheral nerves in ten fetal human hearts (Fig. 8G-L and Supplementary Movies 1-4), as well as markers of other cell types (*eg*. ALDH1A1, FABP4, MYH6, POSTN; Fig. S19). Light-sheet confocal microscopy resolved expression of these proteins at a cell-to-tissue scale, to bridge annotations of cell types and their true spatial distributions.

The great arteries of the outflow tract were more densely innervated than the cardiac apex, and innervation progressed in a proximal-to-distal manner over time between 7 and 10 pcw (Fig. 8G, J; Fig. S19F-H). Nerves tracked with blood vessels on the outside of the heart before penetrating the muscle more robustly in the ventricles than the atria (Fig. 8G, J). The endocardial lining of all valvular cusps and atria, like the arterial trunk and coronary veins, showed sparse PECAM1 immunofluorescence compared to the strong signal of coronary arteries (Fig. S19).

The valves themselves were devoid of nerves at all stages examined (Fig. 8I, L; Fig. S19F). DEGs of FbVIC, NE and EndoCd (Table S1) respectively encoded candidate repulsion mediators annotated by Gene Ontology as mediating negative chemotaxis (GO:0050919) such as *ROBO1*, *SEMA6A*, *SEMA3D* and *PLXNA4* (FbVIC); *SLIT3*, *SEMA3C*, *SEMA6D*, *UNC5B, UNC5*C, *UNC5D*, *DCC* and *DSCAM* in cardiac neurons; and *NTN1*, *PLXNA4* and *PLXND1* in the endocardium. We used CellChat (Jin et al., 2024) to identify the most likely pairs of transcripts encoding ligands from the valves and receptors on axons or their SCP. CellChat predicted that the most significantly probable interaction would be engagement of PTPRZ1 receptors on nerve Schwann cells by inhibitory ligands MDK and PTN (Fukada et al., 2006), produced by endothelial cells and FbVIC (Fig. S17, Fig. S20A). Network analysis associates these three interactants with “positive regulation of glial differentiation (GO:456787)” (Fig. S20B-C), implying that perivalvular nerves may stabilize rather than sprout into leaflets. Nerves themselves could be prevented from pathfinding there by FbVIC-secreted PTPRS or EnTh-produced SEMA3D (Fig. S20A), also likely candidates.

## Discussion

In this work, we independently analyzed whole fetal human hearts using single-cell and spatial transcriptomics, and enriched cell-type annotations with spatial and anatomical information from additional samples, to obtain a comprehensive overview into a vulnerable window of human cardiac development. We highlight an approach also taken by others (Farah et al., 2024; Knight-Schrijver et al., 2022) to conduct iterative refinements based on fully accessible data to update “ground truth” in all developing organ atlases. While new contributions, whether replicative or novel, could seem incremental, they play a critical role in addressing gaps in current knowledge by increasing statistical power. Building a comprehensive, community-driven developmental human cell atlas requires an open, iterative process where scientists can continuously reinterpret and enhance their discoveries in light of new findings and technical approaches (Haniffa et al., 2021).

### Identification of previously undescribed cell types and states

We distinguished nine populations of cardiomyocytes after analyzing 49,227 nuclear transcriptomes from human hearts at 8.6, 9.0, and 10.7 pcw. Among these, two were in a state of preparing for or undergoing active proliferation (CmG2M and CmMit). These comprised 14% and 3% of total cardiomyocytes, respectively, and are at least ten times scarcer in postnatal human hearts (Bergmann et al., 2009). In mice, most cardiomyocytes undergo a final round of nuclear mitosis and cell division near birth, producing post-mitotic, multinucleated or polyploid cells needed to build a sarcomeric scaffold for a lifetime of contraction (Soonpaa et al., 1996). Similarly, most postnatal human heart cells are post-mitotic. This atlas provides a critical reference to normal human development for improving maturation of induced pluripotent stem cells for cell therapy in such conditions as hypoplastic left heart syndrome (Krane et al., 2021) or infarction (Jiang et al., 2024) or to study principles of human cardiac self-organization (Kostina et al., 2024; Lewis-Israeli et al., 2021).

Integration with earlier available datasets revealed a trajectory diversifying novel fibroblast types such as FbVIC from a common progenitor to stromal and cardiomyocyte lineages. Genetic lineage tracing in stem cell-derived organoid or gastruloid models could clarify the potential, contributions and responses of such cardiomyofibroblast progenitors (Argiro et al., 2024; Kostina et al., 2024). Further dataset integration when made available (Bayraktar et al., 2024; Cranley et al., 2024; Lázár et al., 2024) may resolve more of the nature, contributions and competence of stromal cell types in the fetal heart and beyond.

### Stromal diversity reflects anatomical specializations

Fibroblasts are extracellular matrix and growth factor-secreting cells of remarkable functional, organizational and lineage diversity. Epicardial cells predominantly contribute to intracardiac fibroblasts in mice (Ali et al., 2014). However, endocardial and endothelial cells also contribute, particularly in the ventricular septum and chamber walls (Deng et al., 2023; Liu et al., 2018). Furthermore, neural crest cells can differentiate into migrating pericytes, fibroblasts, SMC and even melanocytes in the proximal great vessels, valves, and the septal atrioventricular junction (Etchevers et al., 2001; Hulin et al., 2019; Jiang et al., 2000; Le Lièvre and Le Douarin, 1975). Although no melanocytes were identified in these first-trimester human hearts, *CHD6* (cadherin-6) emerged as a marker of neural crest-derived Class VI cells, with some persistent mesenchymal features at the stages assessed (Clay and Halloran, 2014).

According to the specific valve and even leaflet, valvular interstitial cells (FbVIC) have distinct developmental origins but convergent functions. In atrioventricular (inlet) valves, FbVIC arise mostly from endocardial and epicardial cells, while arterial (outflow) valves include contributions from neural crest, endocardial, and second heart field cells (El Robrini et al., 2016; Etchevers et al., 2001; Jiang et al., 2000; Le Lièvre and Le Douarin, 1975; Liu et al., 2018; Liu et al., 2023; Waldo et al., 1998). FbVIC and SMC are spatially restricted, but Fb1 is also, with more cells in the *tunica adventitia* and proximal ventricles. Our findings confirm that many human cardiac fibroblasts derive from non-epicardial progenitors, as in mice (Deng et al., 2023). Trajectory analysis over integrated 5-10 pcw data shows that epicardial cells had already segregated from the common cardiomyofibroblast (CmF) lineage branch in cardiogenic mesoderm. By 8.6 pcw, CmF had diverged into dividing progenitors contributing preferentially to either stromal or cardiomyocyte lineages. CmF-derived stroma likely contributes via Fb3 fibroblasts to diverse mesenchymal cell types, with FbVIC segregating earlier than other fibroblasts or SMC.

### Complementary information from ST adds atlas precision

Spot clusters, dominated by cells with similar transcriptional profiles, link cardiac cell localization and function when mapped back to anatomically and functionally distinct regions. For instance, *BMP10* was expressed in ventricular trabeculae, as shown in chicken and mouse (Chen et al., 2004; Teichmann and Kessel, 2004), but also in the right atrium, consistent with its asymmetric enrichment in pectinate muscles of mice and humans (Ho et al., 2002; Sedmera et al., 2000). While BMP10’s role remains unclear, thickened pectinate muscle junctions with the lateral right atrial wall of diseased adult human hearts *ex vivo* correlate with persistent atrial fibrillation (Zhao et al., 2023).

This atlas shows that *BMP10* co-varies spatially in the right atrium and in specific cardiomyocytes (CmA1/CmA4 populations; Table S1) with the *PRKAG2* gene (Fig. S21). *PRKAG2* loss-of-function is responsible for congenital, progressive cardiomyopathies with high rates of life-threatening atrial fibrillation and/or ventricular hypertrophy, glycogenosis or tachycardia (Blair et al., 2001; Gollob et al., 2001; White-Brown et al., 2024). A human microdeletion syndrome affecting BMP dosage phenocopies *PRKAG2* hypomorph alleles (Lalani et al., 2009). Our observation provides further evidence that BMP receptor-mediated signal transduction is needed to establish the *annulus fibrosis* at the atrioventricular junction (Gaussin et al., 2005), preventing damaging electrical coupling outside of the pacemaker circuit.

We observed left atrial *PITX2* expression in both transcriptomic modes. Early lateralization reinforced by hemodynamics enhance molecular differences between left and right heart cells in many embryonic models (Hill et al., 2019; Liu et al., 2002; Nowotschin et al., 2006). Laterality establishment by *Pitx2* influences outflow tract morphogenesis and restricts pacemaker development to the right atrium (Hill et al., 2019; Wang et al., 2010). PITX2 remains necessary for adult cardiac adaptive responses (Kahr et al., 2011; Tao et al., 2016).

Integrating all ST data helped us assign spots to sparse structures like the sinoatrial node or an autonomic ganglion. Spot deconvolution with the identities established by our single-nuclear analyses improved prediction of the heterogeneous cellular composition of each spot, and, in return, refined snRNAseq cluster annotation with spatial information. This was true for neuroendocrine, endothelial *versus* endocardial, and trabecular cardiomyocyte identities.

We clarified the historical ambiguity about whether “endocardial” describes a positional or cell-autonomous distinction from endothelial cells of arteries, veins, lymphatics and capillaries (Wang et al., 1998); human fetal endocardium has both spatial and molecular specificities. Furthermore, the multi-modal transcriptomic approach discriminated spatially restricted gene expression, such as *PENK,* within distinct cell types. *PENK* encodes proenkephalin, a component of the neuroendocrine opioid system expressed in murine neural crest-derived mesenchyme and valve primordia (Chen et al., 2022; Liu et al., 2019; Soldatov et al., 2019), was primarily in human FbVIC. (Fig. 6). A skeletal muscle actin we showed to be specifically expressed in maturing *chordae tendineae* is encoded by *ACTA1*, of which variants are associated with dilated cardiomyopathy and defective cardiomyocyte contractility (Garg et al., 2024). The *tunica adventitia*, encasing the proximal ventricles and outflow tract, was best distinguished from other stroma in ST sections. We could more efficiently predict left *versus* right ventricles, compact *versus* trabecular myocardium and conduction nodes *versus* likely differentiating Purkinje cells in subpopulations of the snRNAseq datasets, finding specific and sensitive markers to validate with these complementary techniques.

The combination of snRNAseq and ST provided complementary insights. The former offered quantitative co-transcriptional information at single-nucleus resolution, while the latter provided landmark associations such as chambers or valves, and coordinates relative to the body plan (dorsal/ventral, anterior/posterior) and the organ itself (proximal/distal, inside/outside). This integrative approach based on limited but well-chosen samples can be applied to other organ atlases, adding informative dimensions by leveraging existing and new data.

### Sparse but widespread anatomical networks

The aforementioned methods may bypass rare and/or sparse cells in a sample, including nerves, lymphatics, specialized impulse-conducting cells, resident immune cells, vasculature and fascia. Synergy between next-generation sequencing- or image-driven transcriptomics, high-dimensional protein/mRNA labelling approaches, and qualitative and quantitative measurements enhances functional significance of cellular identity labels. Our 3D immunofluorescence image datasets for a selected time points and markers are a proof-of-concept that multiple networked biological structures can act as anatomical coordinates in the developing fetal and adult organs (Boppana et al., 2023). These datasets can support quantitative segmentation measurements over time and deep-learning model training (Kaltenecker et al., 2024).

Until this year, cell catalogues of the human heart had sampled relatively few individual cells during the critical late first-trimester period, overlooking important cell types and states. As performed for reasons of scale, sampling representative portions of larger organs (Litviňuková et al., 2020) or regressing out highly variable cell-cycle or ribosomal transcripts, as once performed in single-cell analysis pipelines (Butler et al., 2018), can exclude expected and functionally important minority populations. Autonomic neurons and glia, specialized attachment zones between tissues such as the *chordae tendineae*, endothelia, FbVIC and certain cardiomyocyte and mesenchymal progenitor states are among these. Our atlas iteration of the developing human heart situates these populations in their spatiotemporal and functional contexts, providing a reference and roadmap for future research.

## Methods

### Human embryo collection, staging and quality control

First trimester human embryos and fetuses (6-13 pcw) were obtained from induced terminations of pregnancy performed legally in France and sought for reasons other than known fetal abnormality. Tissues were collected after obtaining written consent from donors following procedural initiation, a protocol approved by the national Agency for Biomedical Research (authorization #PFS14-011 to SZ; Agence de la Biomédecine) and declared by the Human Developmental Cell Atlas (HuDeCA; https://hudeca.com) consortium to the French Ministry of Higher Education and Research under reference DC-2022-5011.

### Sample collection

Terminations of pregnancy were induced using a standard combined mifepristone and misoprostol protocol, followed by aspiration. Gestational age was initially determined by ultrasound, refined by measurement of foot length (Evtouchenko et al., 1996; Hern, 1984; O’Rahilly and Müller, 1987; O’Rahilly and Müller, 2010), and confirmed through comparison and contribution of high-resolution photographs of our samples to a staged morphometric and histological cardiac atlas hosted at the Human Developmental Biology Resource (Gerrelli et al., 2015) (https://hdbratlas.org/organ-systems/cardiovascular-system/heart/dissections/gallery.html or https://bit.ly/HumanFetalHeart).

Intact hearts were recovered from aspiration detritus under a Zeiss binocular microscope, rinsed in ice-cold phosphate-buffered saline (PBS) and maintained on ice in Hank’s buffered saline solution until further processing. Retained tissues were documented in a secure HuDeCA-wide database https://hudeca.genouest.org) using the OpenSpecimen biobanking LIMS (https://github.com/krishagni/openspecimen/; Krishagni Solutions, India).

### Genotyping

Quantitative PCR was performed in duplicate on genomic DNA derived from lung or umbilical cord fragments using primers within *SRY* (ACAGTAAAGGCAACGTCCAG and ATCTGCGGGAAGCAAACTGC) to amplify a Y chromosome-specific fragment of 293 bp (Lardenois et al., 2023), and normalized to a 183 bp fragment of the *GAPDH* promoter on chromosome 12, amplified by CCACAGTCCAGTCCTGGGAACC and GAGCTACGTGCGCCCGTAAAA.

Genomic DNA from all samples were further annotated by molecularly determined sex and exclusion of large chromosomal-level copy number variations using a custom digital-droplet PCR analysis (StemGenomics iCS-Digital karyotypes (Assou et al., 2020)). Samples with ambiguous results were reanalyzed using subtelomeric Multiplex Ligation-dependent Probe Amplification (Aneuploidy MLPA probemix [P095], MRC Holland) or for sample HEF_DN_F0000467, with a diagnostic CGH array to exclude a smaller potential copy-number anomaly on chr22 with a high degree of confidence.

### Single-nucleus RNA-sequencing

Three intact hearts from one female and two male fetuses (validated in the transcriptional data) were dissociated for single-nucleus incorporation into the GEMs of the 10X Chromium platform. Dissociation was begun by an initial crush in liquid nitrogen, continued with 10-20 strokes in a Dounce homogenizer over ice, and nuclei further separated from debris over a discontinuous density gradient at 30-35% iodixanol as per a community protocol (Martelotto, 2021). Libraries were prepared by the Genomics and Bioinformatics facility (GBiM) of the U1251 / Marseille Medical Genetics lab using the Chromium Next GEM Single Cell 3= Kit v3.1 (PN-1000269) with Dual Index Kit TT Set A (PN-1000215, 10X Genomics), according to manufacturer’s instructions.

### Sequencing and raw data processing

All libraries were sequenced on a Nextseq 500 (Illumina) by the GBiM facility, except for samples HEF_DN_F0000467 and HEF_DN_F0000530 on a Nextseq 500 by Integragen (Evry, France). The sequenced snRNAseq libraries were processed and aligned to the human reference genome (GRCh38-2020-A) using Cell Ranger software v.3.1.0 (10x Genomics), with unique molecular identifiers (UMI) counted for each barcode.

### Analysis, visualization and integration of snRNAseq datasets

Three datasets of snRNAseq from euploid hearts sequenced on different runs (HEF_DN_F0000530, 8.6 pcw, female; HEF_DN_F0000374, 9.0 pcw, male; HEF_DN_F0000467, 10.7 pcw, male) were analyzed using default parameters in the Seurat 4.1.1 pipeline and 40 PCA dimensions (Hao et al., 2021; Stuart et al., 2019). The pipeline included QC and data filtration, identification of highly variable genes, dimensional reduction, graph-based clustering, and differential expression analysis for the identification of cluster markers. Each dataset was preprocessed and analyzed separately at first, and later integrated with anchor-based batch correction. Further sample details and photographs are in Supplementary Materials and Methods.

### Integration of heart scRNA-seq and snRNA-seq datasets

We integrated our dataset (snRNA-seq) with scRNA-seq datasets of human heart from early stages using the top 1000 HVGs and reciprocal PCA (RPCA) in Seurat; specifically, the datasets included CS12-16 (Xu et al., 2023) and 6.5-7 post-conceptional weeks (Asp et al., 2019). The similarity between cell types across datasets was defined as 1/(1 + distance), where the distance between cell types was determined using a Mehalanobis-like distance metric in Slingshot (Street et al., 2018) based on the top 30 principal components (PCs) of the integrated dataset. Cell types were matched between adjacent developmental stages. The best-matching cell type was initially linked if similarity > 0.06. A secondary match was linked if the similarity z-score across all cell types was > 1 and similarity > 0.06.

### Integration of cardiomyocyte and stromal cells from three datasets

The UMI count matrices of cardiomyocyte and stromal cells from each dataset were combined in Seurat. Integration was performed using the top 1000 HVGs and RPCA, followed by reclustering with top 30 PCs and resolution 0.4. Sixteen clusters were obtained in reclustering for cardiomyocyte and stromal cells across three datasets. PHATE analysis (Moon et al., 2019) was performed on the cardiomyocyte and stromal cells using PCA input and 20 KNNs (k-nearest neighbors). Clusters identified during reclustering were mapped onto the PHATE embedding. The trajectories of those clusters were computed using Slingshot [PmID 29914354] with PCA input.

To classify gene expression along the bifurcation of CmF, we defined four distinct patterns: (1) CmMit > Fb3 and CmF > Fb3; (2) CmMit > CmF and CmMit > Fb3; (3) CmMit < CmF and CmMit < Fb3; (4) CmMit < Fb3 and CmF < Fb3. Here, the ‘<’ and ‘>’ were defined by at least 2-fold change and mean expression > 0.5.

### Spatial transcriptomics

Fresh fetal hearts were staged, photographed under a Leica dissecting microscope and flash-frozen in Leica Frozen Section Compound (FSC)22 in polypropylene molds floated over isopentane cooled by liquid nitrogen. Blocks were stored at -80°C pending quality control (staging, sexing and euploidy). One male heart at 8.4 pcw (HEF_DN_F0000476) and a female heart at 9.7 pcw (HEF_DN_F0000298) were selected. Blocks were trimmed, sectioned at 10 µm and slices melted onto one of: a 10X Genomics Tissue Optimization Slide to determine optimal time for protease digestion (18 minutes); Superfrost Plus (Menzel Gläser) slides for subsequent validation; or a Visium v1 Spatial Gene Expression slide (10X Genomics, one individual heart per slide, with 55 µm spots arrayed at 100 µm center-to-center distance). All were stored with desiccant at -20°C until further processing. As per the Visium v1 Gene Expression protocol, sections were stained in hematoxylin/eosin (H&E), imaged at 10X on an AxioScan 7 (Zeiss, MMG imaging platform), then barcoded mRNA content was captured to generate DNA libraries. Sequencing of libraries was done by the GBiM platform as above.

After standard quality control, reads were aligned to the human reference genome (GRCh38-2020-A) with SpaceRanger v2.0.1 to generate a UMI count matrix per spot. Spots overlaid by tissue were designated from the H&E scans using the manual alignment tool from Loupe Browser v6.4.1 (10X Genomics) to retain only tissue-associated barcodes. Two poorly adherent tissue slices of the 8.4 pcw heart were excluded from further analyses. Two (A90-A91) and four slices (A17-A20) of the younger and older hearts were retained for further analysis and respectively had >3000 median UMI counts per spot under tissue.

#### Analysis, visualization and integration of spatial transcriptomic datasets

Spatial transcriptomic data were first explored by individual section using Loupe v8.0 (10X Genomics) and in further depth with Seurat v5.0.3 (Hao et al., 2024) using 30 dimensions. Clusters were obtained by applying resolutions of 0.45 (section A17), 0.65 (A18), 0.35 (A19), 0.20 (A20), 0.30 (A90) and 0.35 (A91). The R package ‘scTransform’ was used for normalization and variance stabilization (Hafemeister and Satija, 2019). After individual slices were examined and compared, data were integrated using Seurat v5.0.3, spots re-clustered at 0.47 resolution, and clusters manually annotated based on prior knowledge of markers and anatomy.

#### Spatial transcriptomic data deconvolution

RCTD (robust cell-type decomposition (Cable et al., 2022)) from the ‘SpaceX’ (spatially dependent gene co-expression network) R package was used to deconvolute spatial transcriptomic data. Using our integrated snRNAseq data as a reference, we mapped the 21 identified cell types to their spatial distributions on each section. The RCTD “full mode” was used to assign any number of cell types per spot of spatial transcriptomic data. Cell-type weights for each slice, which represent the estimated proportion of each cell type for each spot, were assigned and normalized to sum to 1. This was represented in the form of pie charts for each spot and projected on to the spatial transcriptomic section images to show the estimated proportions of each cell type per spot. CmMit was manually excluded from the snRNAseq reference for deconvolution as it prevented visualization of the true cell type heterogeneity in each spot with strong expression of generic genes as *MYH6* and *MYH7, TOP2A*, *MT-CO1* and *COL1A2*, masking other cell types in the other groups.

### Functional analyses of term annotations

Enrichr-KG (Evangelista et al., 2023) analyses of gene lists were annotated by terms from 2021-2022 releases of the Descartes human fetal atlas (Cao et al., 2020), the Human Gene Atlas (Mabbott et al., 2013), the Tabula Sapiens (The Tabula Sapiens Consortium, 2022) Gene Ontology Biological Processes, and the Kyoto Encyclopedia of Genes and Genomes (KEGG) Human Gene and Pathway databases. Boolean operations on lists of genes were made with GeneVenn (Pirooznia et al., 2007) and/or “Draw Venn Diagrams” (Vandepeer, 2011).

### Histology

#### Hematoxylin and eosin staining

Heart formalin-fixed, paraffin-embedded (FFPE) sections were deparaffinized twice in xylene and progressively rehydrated in ethanol baths of decreasing concentrations diluted in H_2_O. Rehydrated sections were incubated in Mayer’s hematoxylin (Sigma-Aldrich, MHS32) for 2 minutes, rinsed in H_2_O, differentiated briefly in Acid Alcohol (Sigma-Aldrich, A3179), rinsed again for 2 minutes then incubated 5 minutes in aqueous eosin Y (Sigma-Aldrich, HT110232). Sections were dehydrated then mounted with EuKitt (Sigma-Aldrich, 03989). Heart sections were imaged using a Zeiss AxioScan Z1.

### Immunofluorescence

#### Immunofluorescence on cryosection

Hearts were collected in 1X PBS at 4°C and fixed in 4% aqueous buffered paraformaldehyde in PBS (PFA, Electron Microscopy Sciences, 15714) overnight. Hearts were then rinsed in 1X PBS and placed in 15% followed by 30% sucrose in 1X PBS until equilibration at 4°C. Hearts were embedded in OCT (Optimal Cutting Temperature compound, Leica Biosystems, 3801480), oriented, flash-frozen over dry ice or liquid nitrogen and stored at -80°C. Hearts were sectioned to 10 μm thickness using a Leica cryostat. Sections were washed in 1X PBS and fixed 10 minutes in 2% PFA. Sections were permeabilized in 0.1% Triton X-100 (Sigma-Aldrich) in PBS for 20 minutes then incubated in an excess of blocking solution (1% bovine serum albumin, Sigma-Aldrich, A7906; 1% inactivated horse serum, GE Healthcare B15-023; 0.1% Tween-20, Sigma-Aldrich, P1379; in 1X PBS) for 1h. Sections were incubated with primary antibodies diluted in blocking solution overnight at 4°C. After rinsing in PBS, heart sections were incubated with secondary antibodies (Invitrogen, ThermoFisher Scientific) diluted to 1/500 in blocking solution containing 300 nM DAPI (4’,6-diamidino-2-phénylindole) at room temperature for 2 hours. After washes in 1X PBS, slides were mounted in Fluoromount-G (ThermoFisher Scientific, 00-4958-02). Sections were imaged using a Zeiss LSM800 confocal microscope or a Zeiss AxioImager Z2 stereomicroscope with an ApoTome device.

Mouse anti-MYL7 (also known as MLC-2A, Synaptic Systems, 311011) was used at 1/200. Mouse anti-MYH6 (Sigma, AMAb90950) and mouse anti-MYH7 (Abcam ab207926) were diluted to 1/500. Highly cross-absorbed secondary antibodies against relevant species, coupled to AlexaFluor-488, -555 or -647, were all purchased from Thermo Fisher Scientific (refs A11008, A11029, A21244, A31572, A31570, A32787). They were diluted to 1/500 in blocking solution with 300 nM DAPI (4′,6-diamidino-2-phenylindole), and applied to sections for 1h before rinsing.

#### Immunofluorescence on whole hearts

Hearts were collected in cold HBSS, rinsed in PBS and fixed 1 hour in PFA at 4°C with gentle agitation. After washes in 1X PBS, hearts were dehydrated at room temperature 50%, 80% and 100% methanol for 1.5 hour each under agitation. Hearts were then incubated overnight at 4°C in 80% methanol, 6% H_2_O_2_. After progressive rehydration, hearts were incubated in PBSGT (0.2% porcine skin gelatin [VWR 24350.262], 0.5% Triton X-100, 0.01% sodium azide [Sigma-Aldrich, S2002] in 1X PBS) for 3 days with gentle agitation at room temperature. Hearts were then incubated with primary antibodies against CD31/PECAM1 (Cell Signaling Technology, #3528, clone 89C2 at 1/200) and PRPH1 (Sigma-Aldrich AB1530, at 1/1000) at 37°C for 17 to 24 days proportional to relative size, with constant mild agitation. Alternatively, they were incubated in antibodies against ALDH1A1 (Abcam ab52492 clone EP1933Y), S100B (GeneTex GTX14849 clone 4B3), EGR2 (Millipore ABE1374), FABP4 (Bioss BSM-51247M clone 5C1), POSTN (Origene TA804575S clone OTI2B2), or MYH6 (Sigma-Aldrich AMAb90950 clone CL2162). After 24h of washing in six or more changes of PBT (0.5% Triton-X100 in 1X PBS), hearts were incubated with secondary antibodies diluted to 1/1000 (ThermoFisher Scientific, Invitrogen) in PBSGT for 2 days at 37°C. After six or more washes in PBT, hearts were dehydrated. After progressive dehydration in methanol, anhydrous hearts were incubated overnight in brown glass vials with two parts dichloromethane (Sigma-Aldrich, 270997, CAS 75-09-2) to one part methanol until settling, followed by 100% dichloromethane to equilibration. Hearts were next incubated in 100% (di)benzyl ether (Sigma-Aldrich, 108014, CAS 103-50-4) for at least 7 hours (Belle et al., 2017). Transparent hearts were conserved in benzyl ether but transferred to ethyl cinnamate (Sigma-Aldrich, 112372, CAS 103-36-6) for imaging with a Blaze light-sheet UltraMicroscope (Miltenyi Biotec). Imaris software (Oxford Instruments) was used to reconstruct 3D images.

### RNAscope *in situ* hybridization on paraffin sections

Hearts were collected in cold 1X PBS and fixed overnight in 4% PFA. Hearts were then rinsed in 1X PBS and dehydrated in progressive ethanol baths until complete dehydration in 100% ethanol, equilibrated to xylene (twice, 1 hour each), embedded in paraffin and stored at 4°C. 10 μm sections were made using a microtome and mounted on SuperFrost Ultra Plus GOLD slides (Microm Microtech, Epredia, K5800AMNZ72). RNAscope *in situ* hybridization was performed using the RNAscope Multiplex Fluorescent v2 kit from Advanced Cell Diagnostics (ACD), following the manufacturer’s protocol. Before the probe hybridization step, sections were incubated in the “protease PLUS” solution for 15 minutes at 40°C. Up to three of *ISL1* (478591-C3), *PRRX1* (511001-C2), *PENK* (548301-C3), *MEOX1* (564321), *ACTA1* (454311) and *BMP10* (459641) RNAscope probes were then hybridized. Nuclei were counterstained with the provided DAPI solution. Opal 520, Opal 570 and/or Opal 690 reagent packs (Akoya Biosciences) were used at 1/200 dilution to localize probes. Heart sections were imaged with tiling on a Zeiss LSM800 confocal microscope and stitched with Zen 3.7 software.

## Websites

Loupe Browser

https://www.10xgenomics.com/products/loupe-browser/downloads/eula

Seurat vignettes and use-cases

https://satijalab.org/seurat/articles/pbmc3k_tutorial

https://satijalab.org/seurat/articles/sctransform_vignette

https://satijalab.org/seurat/articles/spatial_vignette

Enrichr-KG

https://maayanlab.cloud/enrichr-kg

Venn diagrams and subsets of co-expression

https://www.bioinformatics.org/gvenn/

https://bioinformatics.psb.ugent.be/webtools/Venn/

Human Developmental Cell Atlas (France)

https://hudeca.com/

https://hudeca.genouest.org/

## Acknowledgments

Prof. Aubert Agostini and staff of the Centre de Gynécologie Sociale de l’AP-HM in Marseille obtained patient consents and provided most of the tissue donations used in this work, to which effort Maryne Toupin also contributed from Rennes. Laurence Noël built and helped curate the ontologies used in HuDeCA’s OpenSpecimen database. The authors gratefully acknowledge personnel of MMG’s GBiM and imaging platforms, especially Catherine Robert, Christel Castro, Dr. Valérie Delague, Nicolas Lenfant, Alexandre Atkinson, Deniz Harcanoğlu and Dr. Raphaël Blain for their analysis assistance.

## Funding

Institut National de la Santé et de la Recherche Médicale (INSERM) Programme Transversale “HuDeCA”, France (MH, YG, AC, SMG, SZ, HCE)

Association Française contre les Myopathies grant “MoThARD” (SK, AB, SZ, HCE)

CDB is supported by a grant from the Fondation pour la Recherche Médicale, France.

*This publication is part of the Human Cell Atlas.* https://www.humancellatlas.org/publications/

## Author contributions

SZ and HCE conceptualized the study and designed the experiments. CDB, SZ and HCE refined methodology, interpreted the data and wrote the original manuscript draft, to which AB and SK also contributed. MH, MM, WS, CM and SMG processed samples and performed quality control measures. MH carried out the histological, immunofluorescence and high-throughput sequencing-based experiments. CDB, CH, SK, SR, YX, AB and HCE designed, refined and analyzed bioinformatics experiments. AL, IV, YG and AC contributed to the 3D immunofluorescence experimental design, execution and data analyses. CDB, ZB, IV, AC, AB, SZ and HCE supervised the analyses. All authors reviewed and provided input on the final manuscript.

## Data and materials availability

All data, code, and materials used in the analyses are available either in the main text and/or the Supplementary Materials, or deposited in the following repositories: Sequencing data, images and snRNAseq matrices are available from the Gene Expression Omnibus (GEO), https://www.ncbi.nlm.nih.gov/geo/ under accession GSE283967; code from https://github.com/BAUDOTlab/Human_fetal_heart_atlas; movies, large images and high-resolution gross anatomy images are available from Figshare at https://figshare.com/projects/Multi-modal_refinement_of_the_human_heart_atlas_during_the_first_gestational_trimester/213151. Some images are reproduced at the Human Developmental Biology Resource (UK) fetal cardiac anatomy website (https://bit.ly/HumanFetalHeart).

Human Cell Atlas data from (Xu et al., 2023) is available both at: https://explore.data.humancellatlas.org/projects/e255b1c6-1143-4fa6-83a8-528f15b41038 and from GEO under accession GSE157329.

