## Supplementary material for "Multi-modal refinement of the human heart atlas during the first gestational trimester": Fig. S

### Supplementary data

#### Supplementary figure legends

##### Supplementary figure S1: Histological analysis of human fetal hearts

(A-I) Hematoxylin and eosin staining on different axial level sections of human fetal heart at 8.7 pcw (A-C), 9.7 pcw (D-F) and 11.0 pcw (G-I). RA, right atrium; LA, left atrium; Ao, aorta; PT, pulmonary trunk; CA, coronary artery; AoV, aortic valve; RV, right ventricle; LV, left ventricle; S, interventricular septum; sb, subvalvular domain (septal bridge); lAV, left atrioventricular valve; rAV, right atrioventricular valve; PV, pulmonary valve.

##### Supplementary figure S2: Cross-referencing annotations improves identity assignment

(**A**) Co-expression of *KCNQ5* and *INPP4B* in a UMAP plot of integrated snRNA-seq data of all nuclei from 8.4, 9.0 and 10.7 pcw hearts, showing enrichment of cells transcribing both genes in the CmG2M and CmV1 subpopulations of Class I cells (cardiomyocytes). (**B**) Enrichr-KG analysis (Evangelista et al., 2023) of Class VI clusters converges for cluster 18 on Schwann cells, glia and axon guidance, leading us to annotate it as Schwann cells and their precursors, while in (**C**) analysis of cluster 19 with the same parameters highlights different visceral neuron and synapse annotations. Combined with the knowledge that neuronal nuclei at the level of cardiac ganglia are parasympathetic, we annotated this cluster as "neuroendocrine" (as it also expressed *CHGA*, *CHGB*, *SST* and the Descartes term "chromaffin cells in Adrenal", a neural crest endocrine derivative, was significantly enriched [$p = 3.43 \times 10^{-14}$]).

##### Supplementary figure S3: Determination of similarities across annotated datasets

(**A**) Similarity between cell types across datasets, calculated as 1/(1 + distance). The distance between cell types was determined using the Mehalanobis-like distance metric in Slingshot (Street et al., 2018) based on the top 30 principal components (PCs) of the integrated dataset. Color bars indicate cell type classes. (**B**) Cell type mapping across datasets based on the similarity in panel A. Cell types were matched between adjacent developmental stages. Initially, the best-matching cell type was linked if similarity > 0.06. A secondary match was linked if the similarity z-score across all cell types was > 1 and similarity > 0.06. (**C**) PHATE embedding of the integrated dataset, with cells colored by developmental stages. (**D**) Cells from Asp et al., 2019 in the PHATE embedding of our integrated dataset, showing cell types as originally annotated.

#### Supplementary figure S4: Spatial transcriptomics analysis (ST) of 8.4 pcw A90 heart section

(**A**) Seurat UMAP plot of spatial transcriptomic data from 8.4 pcw A90 heart section. (**B**) Spatial dimensionality plot displaying spatial distribution and cluster annotation. (**C**) Spatial plots showing spatial gene expression, with highest expression in red and lowest expression in blue. Example of marker genes of the myocardium, atrial and ventricular myocardium, trabeculae, aorta and valve are displayed. RV, right ventricle; LV, left ventricle; RA, right atrium; LA, left atrium; Ao, aorta.

#### Supplementary figure S5: ST of 8.4 pcw A91 heart section

(**A**) Seurat UMAP plot of spatial transcriptomic data from 8.4 pcw A91 heart section. (**B**) Spatial dimensionality plot displaying spatial distribution and annotation of clusters. (**C**) Spatial plots showing spatial gene expression, with highest expression in red and lowest expression in blue. Example of marker genes of the myocardium, atrial and ventricular myocardium, trabeculae, aorta and valve are displayed. RV, right ventricle; LV, left ventricle; RA, right atrium; LA, left atrium; Ao, aorta.

#### Supplementary figure S6: ST of 9.7 pcw A17 heart section

(**A**) Seurat UMAP plot of spatial transcriptomic data from 9.7 pcw A17 heart section. (**B**) Spatial dimensionality plot displaying spatial distribution and annotation of clusters. (**C**) Spatial plots showing spatial gene expression, with highest expression in red and lowest expression in blue. Example of marker genes of the myocardium, compact myocardium, atrial and ventricular myocardium, trabeculae, aorta and valve are shown. RV, right ventricle; LV, left ventricle; RA, right atrium; LA, left atrium; Ao, aorta.

#### Supplementary figure S7: ST of 9.7 pcw A18 heart section

(**A**) Seurat UMAP plot of spatial transcriptomic data from 9. 7 pcw heart section number 2. (**B**) Spatial dimensionality plot displaying spatial distribution and annotation of clusters. (**C**) Spatial plots showing spatial gene expression, with highest expression in red and lowest expression in blue. Example of marker genes of the myocardium, compact myocardium, atrial and ventricular myocardium, trabeculae, aorta and valve are shown. RV, right ventricle; LV, left ventricle; RA, right atrium; LA, left atrium; Ao, aorta.

#### Supplementary figure S8: ST of 9.7 pcw A19 heart section

(**A**) Seurat UMAP plot of spatial transcriptomic data from 9.7 pcw heart section number 3. (**B**) Spatial dimensionality plot displaying spatial distribution and annotation of clusters. (**C**) Spatial plots showing spatial gene expression, with highest expression in red and lowest expression in blue.

Example of marker genes of the myocardium, atrial and ventricular myocardium, trabeculae, aorta and valve are shown. RV, right ventricle; LV, left ventricle; RA, right atrium; LA, left atrium; Ao, aorta.

#### Supplementary figure S9: ST of 9.7 pcw A20 heart section

(**A**) Seurat UMAP plot of spatial transcriptomic data from 9.7 pcw heart section number 4. (**B**) Spatial dimensionality plot displaying spatial distribution and cluster annotation. (**C**) Spatial plots showing spatial gene expression, with highest expression in red and lowest expression in blue. Example of marker genes of the myocardium, atrial and ventricular myocardium, trabeculae, aorta and valve are displayed. RV, right ventricle; LV, left ventricle; RA, right atrium; LA, left atrium; Ao, aorta.

#### Supplementary figure S10: Split UMAP plots of integrated spatial transcriptomic data, from two 8.4 pcw and four 9.7 pcw sections with cluster annotations.

Split UMAP plots of the integrated spatial transcriptomic data in Figure 3 according to section, with cluster annotations on the right.

#### Supplementary figure S11: Deconvolution of 8.4 pcw A90 heart section spatial transcriptomic data with integrated snRNAseq data as reference.

(**A**) Deconvolution of spatial transcriptomic data from 8.4 pcw A90 heart section with RCTD software. The proportion of each cell type identified in integrated snRNAseq data (Figure 1) is estimated and represented in pie charts for each spatial transcriptomic pixel. (**B**) Estimated cell type proportion for each snRNAseq cluster on each spatial pixel of 8.4 pcw A90 heart section, with highest estimated proportion in red and lower in blue. RA, right atrium; LA, left atrium; RV, right ventricle; LV, left ventricle.

#### Supplementary figure S12: Deconvolution of 8.4 pcw A91 heart section spatial transcriptomic data with integrated snRNAseq data as reference.

(**A**) Deconvolution of spatial transcriptomic data from 8.4 pcw A91 heart section with RCTD software. The proportion of each cell type identified in integrated snRNAseq data (Figure 1) is estimated and represented in pie charts for each spatial transcriptomic pixel. (**B**) Estimated cell type proportion for each snRNAseq cluster on each spatial pixel of 8.4 pcw A91 heart section, with highest estimated proportion in red and lower in blue. RA, right atrium; LA, left atrium; RV, right ventricle; LV, left ventricle.

#### Supplementary figure S13: Deconvolution of 9.7 pcw A17 heart section spatial transcriptomic data with integrated snRNAseq data as reference.

(**A**) Deconvolution of spatial transcriptomic data from 9.7 pcw heart section number 1 with RCTD software. The proportion of each cell type identified in integrated snRNAseq data (Figure 1) is estimated and represented in pie charts for each spatial transcriptomic pixel. (**B**) Estimated cell type proportion for each snRNAseq cluster on each spatial pixel of 9.7 pcw A17 heart section, with highest estimated proportion in red and lower in blue. RA, right atrium; LA, left atrium; RV, right ventricle; LV, left ventricle.

#### Supplementary figure S14: Deconvolution of 9.7 pcw A18 heart section spatial transcriptomic data with integrated snRNAseq data as reference.

**(A)** Deconvolution of spatial transcriptomic data from 9.7 pcw heart section number 2 with RCTD software. The proportion of each cell type identified in integrated snRNAseq data (Figure 1) is estimated and represented in pie charts for each spatial transcriptomic pixel. **(B)** Estimated cell type proportion for each snRNAseq cluster on each spatial pixel of 9.7 pcw A18 heart section, with highest estimated proportion in red and lower in blue. RA, right atrium; LA, left atrium; RV, right ventricle; LV, left ventricle.

#### Supplementary figure S15: Deconvolution of 9.7 pcw A19 heart section spatial transcriptomic data with integrated snRNAseq data as reference.

**(A)** Deconvolution of spatial transcriptomic data from 9.7 pcw heart section number 3 with RCTD software. The proportion of each cell type identified in integrated snRNAseq data (Figure 1) is estimated and represented in pie charts for each spatial transcriptomic pixel. **(B)** Estimated cell type proportion for each snRNAseq cluster on each spatial pixel of 9.7 pcw A19 heart section, with highest estimated proportion in red and lower in blue. RA, right atrium; LA, left atrium; RV, right ventricle; LV, left ventricle.

#### Supplementary figure S16: Deconvolution of 9.7 pcw A20 heart section spatial transcriptomic data with integrated snRNAseq data as reference.

**(A)** Deconvolution of spatial transcriptomic data from 9.7 pcw heart section number 4 with RCTD software. The proportion of each cell type identified in integrated snRNAseq data (Figure 1) is estimated and represented in pie charts for each spatial transcriptomic pixel. **(B)** Estimated cell type proportion for each snRNAseq cluster on each spatial pixel of 9.7 pcw A20 heart section, with highest estimated proportion in red and lower in blue. RA, right atrium; LA, left atrium; RV, right ventricle; LV, left ventricle.

#### Supplementary figure S17: Valve markers in spatial transcriptomic sections A91 and A17.

Spatial transcriptomic plots showing expression of valve marker genes on 8.4 pcw A91 heart **(A-G)** and 9.7 pcw A17 **(H-K)** sections, with highest expression in red and lowest in blue.

#### Supplementary figure S18: Parasympathetic ganglionic marker gene detection in spatial transcriptomic section A90.

Spatial transcriptomic plots showing expression of parasympathetic ganglia marker genes in the 8.4 pcw A90 heart section, with highest expression in red and lowest in blue.

#### Supplementary figure S19: Use-cases of 3D protein expression data

Segmentation of right and left coronary arteries **(A, B)** from 3D-reconstructed stacks of a 10.1 pcw female fetal heart (F0570) after anti-PECAM1 immunofluorescence, tissue clearing and light-sheet confocal microscopy, shown in **(C)** dorsal and **(D)** ventral views. Ventricles are heavily vascularized with strongly PECAM1-immunoreactive vessels, in particular the coronary arteries (arrows). **(E)** One confocal section of same image stack showing coronary arteries in section at the top of the ventricles and in the ventricular sulcus near the apex, arrowheads. Inset, ventral view of same heart on retrieval, showing the visual difficulty of extrapolating volumes from one or a few sections, but also the difficulty of resolving structures such as valves or endocardial surfaces within the coronary cages in C or D. Also cf. Movies S3 and S4. **(F) Left:** PRPH immunofluorescence at 6.5-7 pcw in a whole heart at approximately 6.5-7 pcw (EE3383). Innervation progresses along the great arteries, circumnavigates the atrioventricular junctions above the coronary sinus and converges at the level of the atrioventricular node at the base of the right atrium before extending distally along the ventricles toward apex (arrow). Right: A different sample at 7 pcw, immunostained to show global PECAM1+ vascular endothelial (cyan) and TNNT1+ myocardial (magenta) distribution. Note absence of atrial innervation. (**G**) PRPH+ immunofluorescent nerves, having attained the apex (diagonal arrow) and extending along distal ventricular edges by 9.1 pcw. Note that only proximal right atrium and most of the left atrium is innervated at this stage. (H) By 10.1 pcw, the great arteries are heavily innervated and PRPH+ fascicles have formed near and around the coronary artery branches, largely sparing the left septal atrium at this stage. The right ventricular nerves have attained the apex (arrow). This view is from the same speciment and perspective as in (**D**). **(I)** ALDH1A1 is strongly immunofluorescent in the coronary and pericardiophrenic arteries (PCPh), in this 7 pcw sample (F0004) with intact pericardium. (**J**) Sample F0004 was also immunolabeled to show weak FABP4 fluorescence in endocardial and coronary vein endothelium (pink). Arrowheads delimit the left ventricle in this side view, dorsal to right. PCw, parietal pericardial wall. (**K**) POSTN is a mesenchymal marker strongly immunofluorescent at the atrioventricular valve hinge points (arrows) above the *annulus fibrosus* among other tissues, including the tracheal rings (t). (**L**) The same heart was simultaneously stained with an antibody against MYH6 strongly labels the atria in this dorsal view of sample F0010 at 8.0 pcw. LA, left atrium; LV, left ventricle; RA, right atrium; RV, right ventricle. Bar = 500 µm in (**A**-**H, K, L)**; in (**I, J**) = 300 µm.

#### Supplementary figure S20: *In silico* interaction and annotation analyses

(**A**) CellChat (Jin et al., 2024) enabled a query of the significance of the likely of interactions mediated by secreted ligands encoded by genes expressed by a first cell type and cognate receptors potentially expressed by an adjacent second cell type. Default settings were used and excluded potential communications between fewer than ten cells. We used this tool to identify and weight the probability of communication between either valvular FbVIC or EnTh cells and either SCP or NE on autonomic nerves at the base of valvular leaflets. FbVIC are predicted to be highly likely to secrete PTN, received by PTPRZ1 on SCP, as well as PTPRS, PTN and IGF1 to potentially act on NTRK3, ALK and IGF1R, respectively, on NE. A selection of interactions is represented here, and the most likely in red, based on expression levels and proximity, are circled. (**B**) Enrichr-KG analysis of a selection of potential molecules to mediate axon guidance highlights the Gene Ontology annotations often shared between them, with the PTPRZ1-PTN-MDK triad associated with terms related to "positive regulation of glial cell differentiation". *SEMA3D* and *SEMA3A* are associated with "negative regulation of axon extension involved in axon guidance". (C) The top twenty highly significantly associated GO Biological_Process_2021 terms are headed by "axon guidance (GO:0007411)" and contain "regulation of axon extension involved in axon guidance (GO:0048841)" which annotates *SEMA3A*, *SEMA3D* and *NRP1*, among other genes.

#### Supplementary figure S21: ST plots of *PRKAG2*

*PRKAG2* is relatively more expressed in atrial cardiomyocytes than in ventricular populations, as seen in sections (A) A90, (B) A91 at 8.4 pcw and (C) A17 and (D) A19, at 9.7 pcw. Cf. Table S1. By the later stage, there is preponderant expression in the right atrium.

### Supplementary Movies

#### Movie 1.

Heart F0495, 9.1 pcw male; immunofluorescence of PECAM1 (pink, vascular endothelium) and PRPH (yellow, peripheral nerves) followed by modified iDISCO tissue clearing. Reconstruction of signal across 1288 optical sections acquired on a Miltenyi Blaze light-sheet confocal microscope using Imaris software.

#### Movie 2.

Heart F0514, 9.1 pcw female; immunofluorescence of PECAM1 (pink, vascular endothelium) and PRPH (yellow, peripheral nerves) followed by modified iDISCO tissue clearing. Reconstruction of signal across 1332 optical sections acquired on a Miltenyi Blaze light-sheet confocal microscope using Imaris software.

#### Movie 3.

Heart F0570, 10.1 pcw female; immunofluorescence of PECAM1 (pink, vascular endothelium) and PRPH (yellow, peripheral nerves) followed by modified iDISCO tissue clearing. Reconstruction of signal across 1340 optical sections acquired on a Miltenyi Blaze light-sheet confocal microscope using Imaris software.

#### Movie 4.

Heart F0570, 10.1 pcw female; immunofluorescence of PECAM1 (pink, vascular endothelium) and PRPH (yellow, peripheral nerves) followed by modified iDISCO tissue clearing. Segmentation by clipping planes in Imaris software to show PECAM1-stained cusps of aortic valve and departure of left and right coronary arteries.

### Supplementary table legends

#### Supplementary Materials and Methods

Summary of samples, stages, unique identifiers when available, analyses conducted and information about transcriptomics per sample in the summary tab. Photos of whole hearts, when available, are linked from the HuDeCA ID to the gross anatomy tab.

#### Supplementary Table 1

Data pertaining to 49,227 nuclear transcriptomes from three first-trimester human hearts. **First tab**: Index with hyperlinks to other tabs in table. Within all other tabs, a hyperlinked “Index” button will return to this tab. **Second tab**: Cluster names and cell statistics, broken down by source material. **Third tab**: All genes differentially expressed per cluster, relative to all others. **Fourth tab**: as previous but featuring top 20 differentially expressed genes per cluster, relative to all others. **Subsequent tabs**: All genes differentially expressed per class or cluster, relative to all other classes or to other cluster(s) in same tab. Abbreviations: *p_val*, p-value; *avg_log2FC*, average log(2) fold-change; *pct.1*, percentage of cluster cells where the gene is detected; *pct.2*, percentage of all other cells where the gene is detected; *p_val_adj*, p-values adjusted for multiple hypothesis testing (based on a Bonferroni correction); *Cluster.no*, local snRNAseq cluster number; *gene*, official human gene name.

#### Supplementary Table 2

Marker genes and statistics of clusters from integrated snRNA-seq and scRNA-seq data, referring to Figure 2.

#### Supplementary Table 3

Marker genes and statistics of clusters from individual spatial transcriptomic analyses of A90, A91, A17, A18, A19 and A20 heart section data.

#### Supplementary Table 4

Marker genes and statistics of spatial transcriptomic clusters integrated from section A90, A91, A17, A18, A19 and A20 data.

#### Supplementary Table 5

Average proportions of each cell type from integrated snRNAseq data within each spatial transcriptomic cluster after deconvolution, per section.

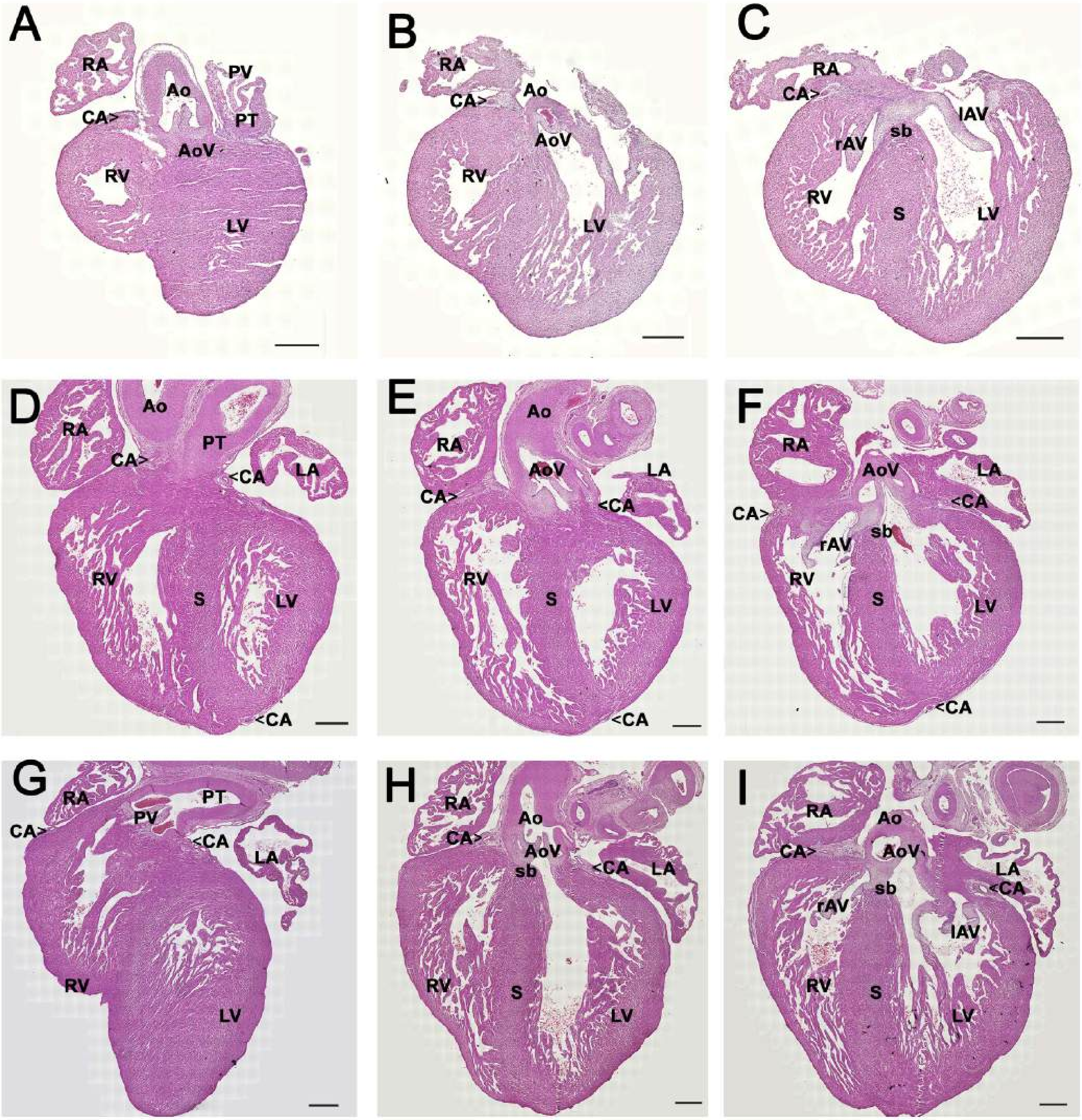
A
RA
Ao
PV
PT
CA>
AoV
RV
LV
B
RA
CA>
Ao
AoV
RV
LV
C
RA
CA>
lAV
rAV
sb
RV
S
LV
D
Ao
RA
PT
CA>
<CA
LA
RV
S
LV
<CA
E
Ao
RA
AoV
LA
CA>
<CA
RV
S
LV
<CA
F
RA
AoV
LA
<CA
CA>
sb
rAV
RV
S
LV
<CA
G
RA
PT
CA>
PV
<CA
LA
RV
LV
H
RA
Ao
CA>
AoV
sb
<CA
LA
RV
S
LV
I
RA
Ao
CA>
AoV
LA
sb
<CA
rAV
lAV
RV
S
LV

Fig S1

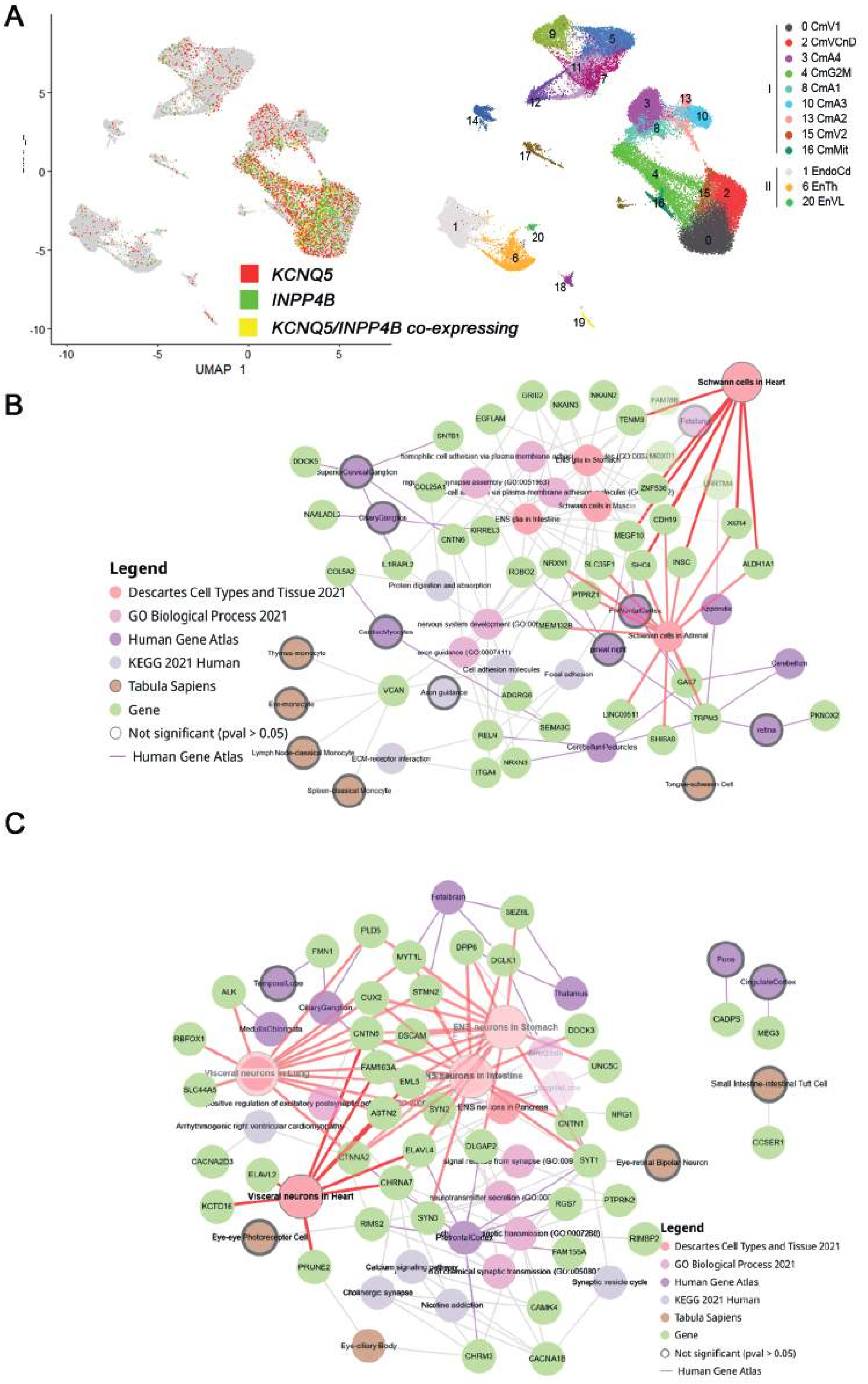
A
KCNQ5
INPP4B
KCNQ5/INPP4B co-expressing
UMAP 1
0 CmV1
2 CmVCnD
3 CmA4
4 CmG2M
8 CmA1
10 CmA3
13 CmA2
15 CmV2
16 CmMit
1 EndoCd
6 EnTh
20 EnVL
5 Fb1
7 Fb3
9 FbVIC
11 Fb2
12 SMC
14 EpiCd
17 Bl
18 SCP
19 NE
B
Schwann cells in Heart
Schwann cells in Adrenal
ENS glia in Stomach
ENS glia in Intestine
Schwann cells in Muscle
Legend
Descartes Cell Types and Tissue 2021
GO Biological Process 2021
Human Gene Atlas
KEGG 2021 Human
Tabula Sapiens
Gene
Not significant (pval > 0.05)
Human Gene Atlas
C
ENS neurons in Stomach
ENS neurons in Intestine
ENS neurons in Pancreas
Visceral neurons in Lung
Visceral neurons in Heart
Arrhythmogenic right ventricular cardiomyopathy
Cholinergic synapse
Nicotine addiction
Synaptic vesicle cycle
Small Intestine-intestinal Tuft Cell
Legend
Descartes Cell Types and Tissue 2021
GO Biological Process 2021
Human Gene Atlas
KEGG 2021 Human
Tabula Sapiens
Gene
Not significant (pval > 0.05)
Human Gene Atlas

Fig S2

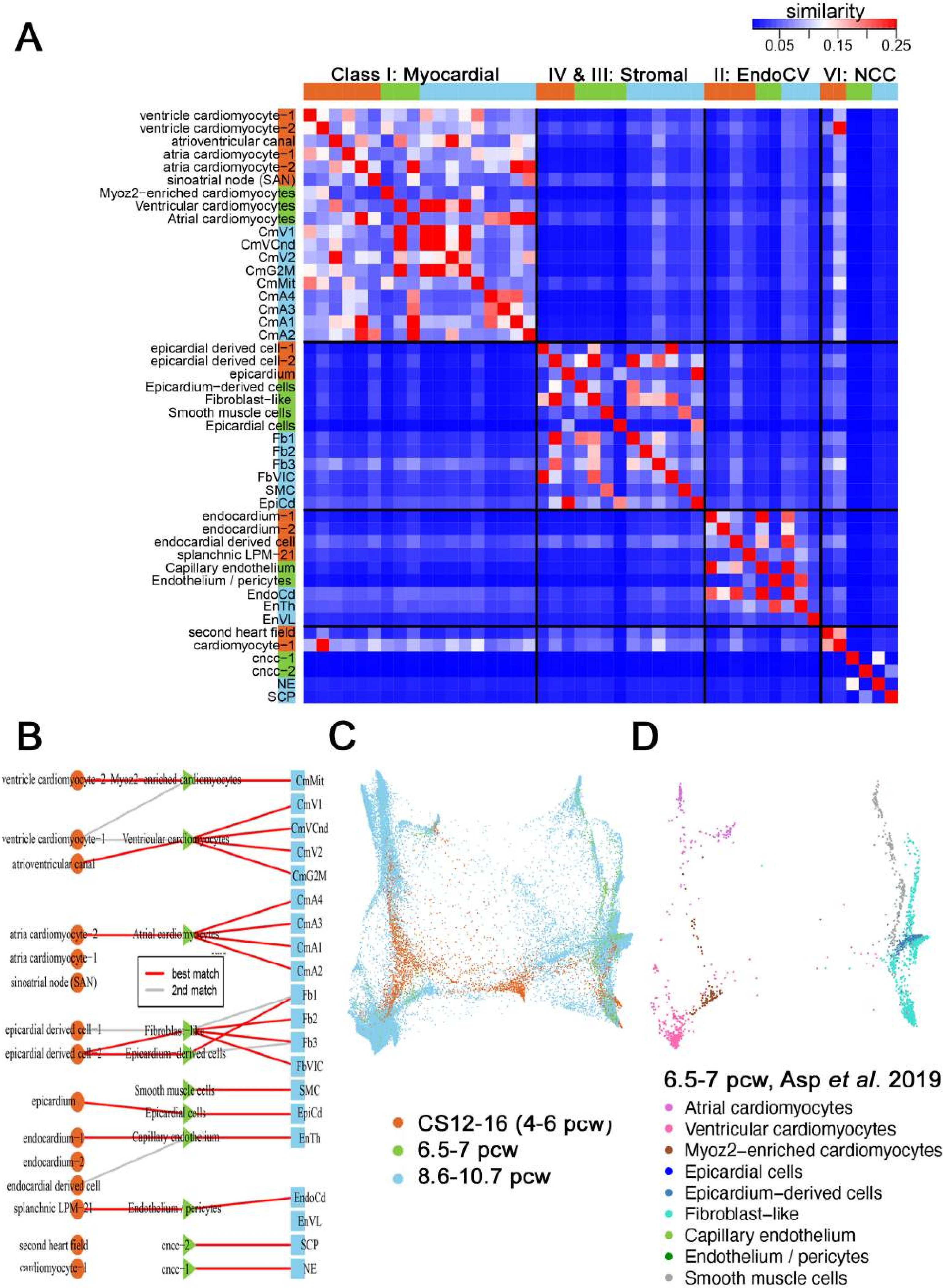

A
similarity
0.05
0.15
0.25
Class I: Myocardial
IV & III: Stromal
II: EndoCV
VI: NCC
ventricle cardiomyocyte−1
ventricle cardiomyocyte−2
atrioventricular canal
atria cardiomyocyte−1
atria cardiomyocyte−2
sinoatrial node (SAN)
Myoz2−enriched cardiomyocytes
Ventricular cardiomyocytes
Atrial cardiomyocytes
CmV1
CmVCnd
CmV2
CmG2M
CmMit
CmA4
CmA3
CmA1
CmA2
epicardial derived cell−1
epicardial derived cell−2
epicardium
Epicardium−derived cells
Fibroblast−like
Smooth muscle cells
Epicardial cells
Fb1
Fb2
Fb3
FbVIC
SMC
EpiCd
endocardium−1
endocardium−2
endocardial derived cell
splanchnic LPM−21
Capillary endothelium
Endothelium / pericytes
EndoCd
EnTh
EnVL
second heart field
cardiomyocyte−1
cncc−1
cncc−2
NE
SCP
B
best match
2nd match
C
CS12-16 (4-6 pcw)
6.5-7 pcw
8.6-10.7 pcw
D
6.5-7 pcw, Asp *et al.* 2019
Atrial cardiomyocytes
Ventricular cardiomyocytes
Myoz2−enriched cardiomyocytes
Epicardial cells
Epicardium−derived cells
Fibroblast−like
Capillary endothelium
Endothelium / pericytes
Smooth muscle cells


Fig S3

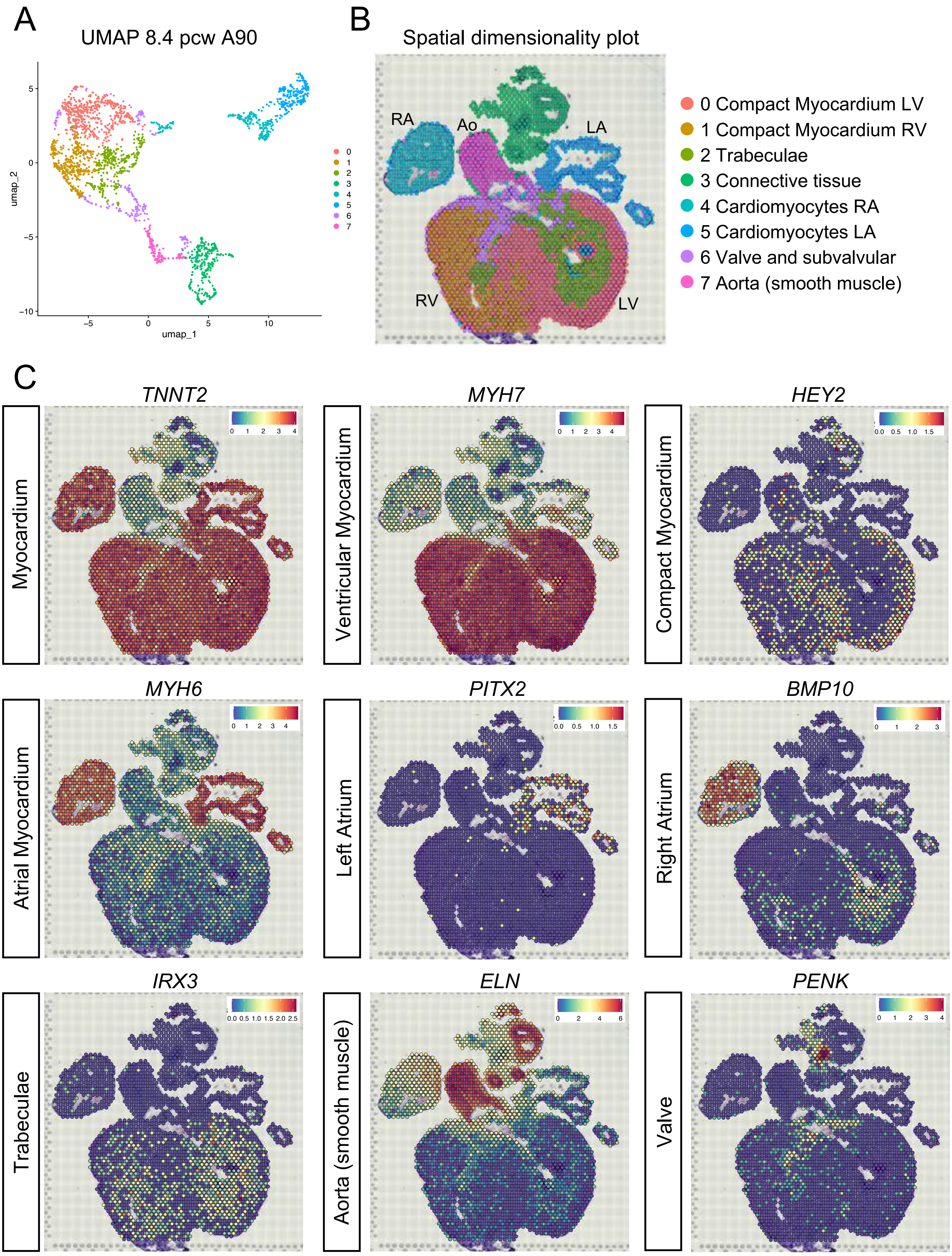
A
UMAP 8.4 pcw A90
umap_2
umap_1
B
Spatial dimensionality plot
RA
Ao
LA
RV
LV
0 Compact Myocardium LV
1 Compact Myocardium RV
2 Trabeculae
3 Connective tissue
4 Cardiomyocytes RA
5 Cardiomyocytes LA
6 Valve and subvalvular
7 Aorta (smooth muscle)
C
Myocardium
TNNT2
Ventricular Myocardium
MYH7
Compact Myocardium
HEY2
Atrial Myocardium
MYH6
Left Atrium
PITX2
Right Atrium
BMP10
Trabeculae
IRX3
Aorta (smooth muscle)
ELN
Valve
PENK

Fig S4

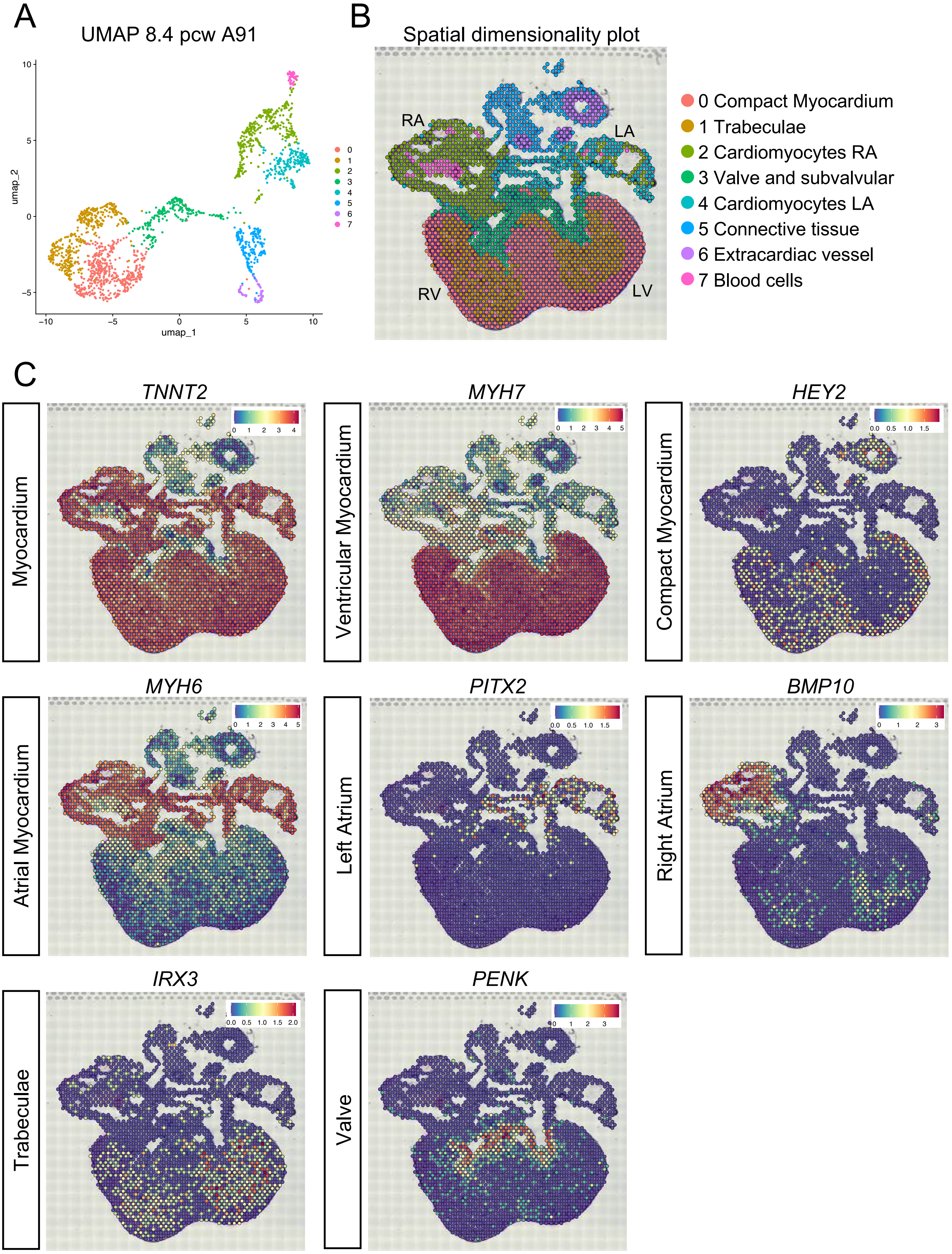
A
UMAP 8.4 pcw A91
umap_2
umap_1
B
Spatial dimensionality plot
RA
LA
RV
LV
0 Compact Myocardium
1 Trabeculae
2 Cardiomyocytes RA
3 Valve and subvalvular
4 Cardiomyocytes LA
5 Connective tissue
6 Extracardiac vessel
7 Blood cells
C
Myocardium
TNNT2
Ventricular Myocardium
MYH7
Compact Myocardium
HEY2
Atrial Myocardium
MYH6
Left Atrium
PITX2
Right Atrium
BMP10
Trabeculae
IRX3
Valve
PENK

Fig S5

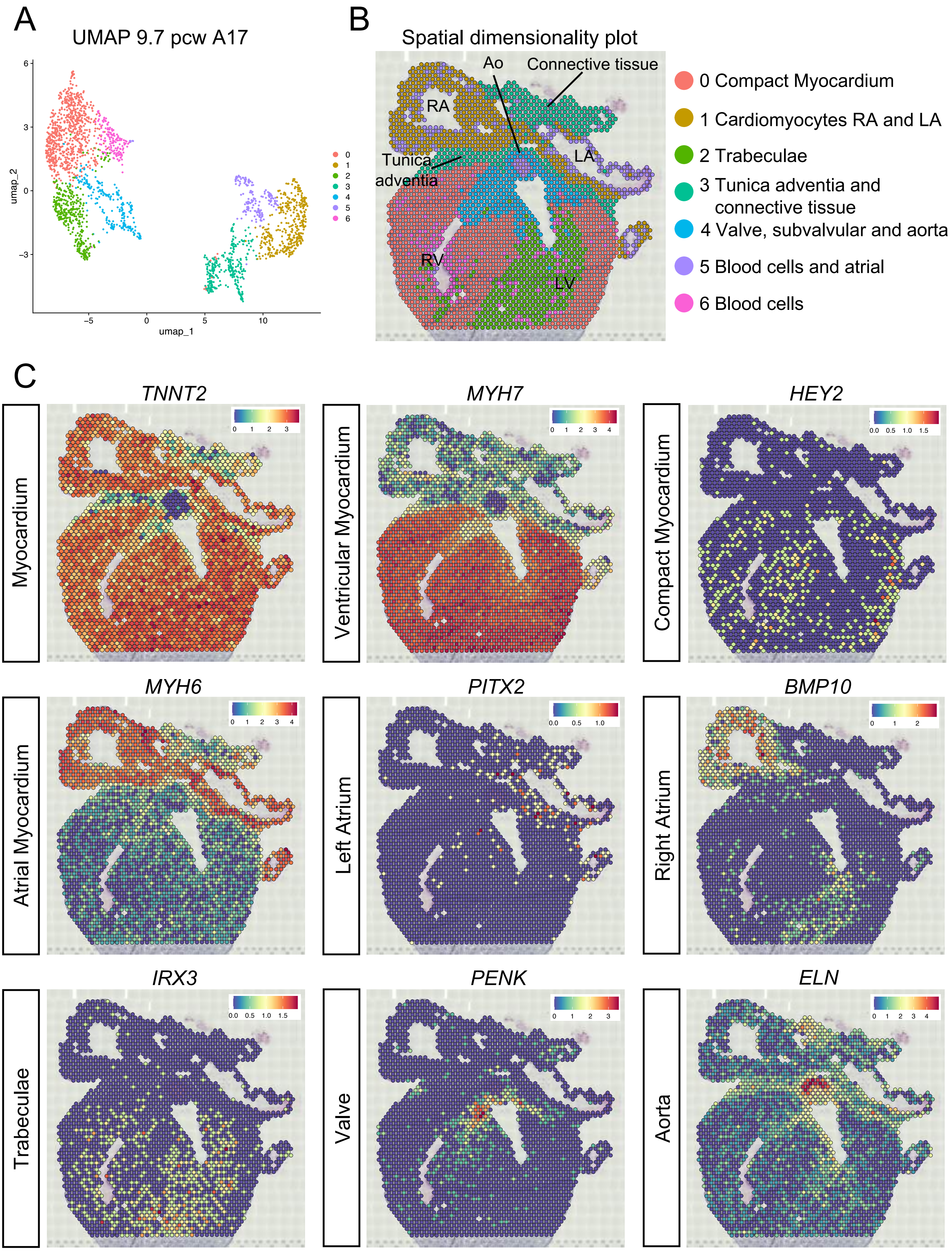
A
UMAP 9.7 pcw A17
umap_2
umap_1
B
Spatial dimensionality plot
Ao
Connective tissue
RA
Tunica adventia
LA
RV
LV
0 Compact Myocardium
1 Cardiomyocytes RA and LA
2 Trabeculae
3 Tunica adventia and connective tissue
4 Valve, subvalvular and aorta
5 Blood cells and atrial
6 Blood cells
C
TNNT2
MYH7
HEY2
Myocardium
Ventricular Myocardium
Compact Myocardium
MYH6
PITX2
BMP10
Atrial Myocardium
Left Atrium
Right Atrium
IRX3
PENK
ELN
Trabeculae
Valve
Aorta

Fig S6

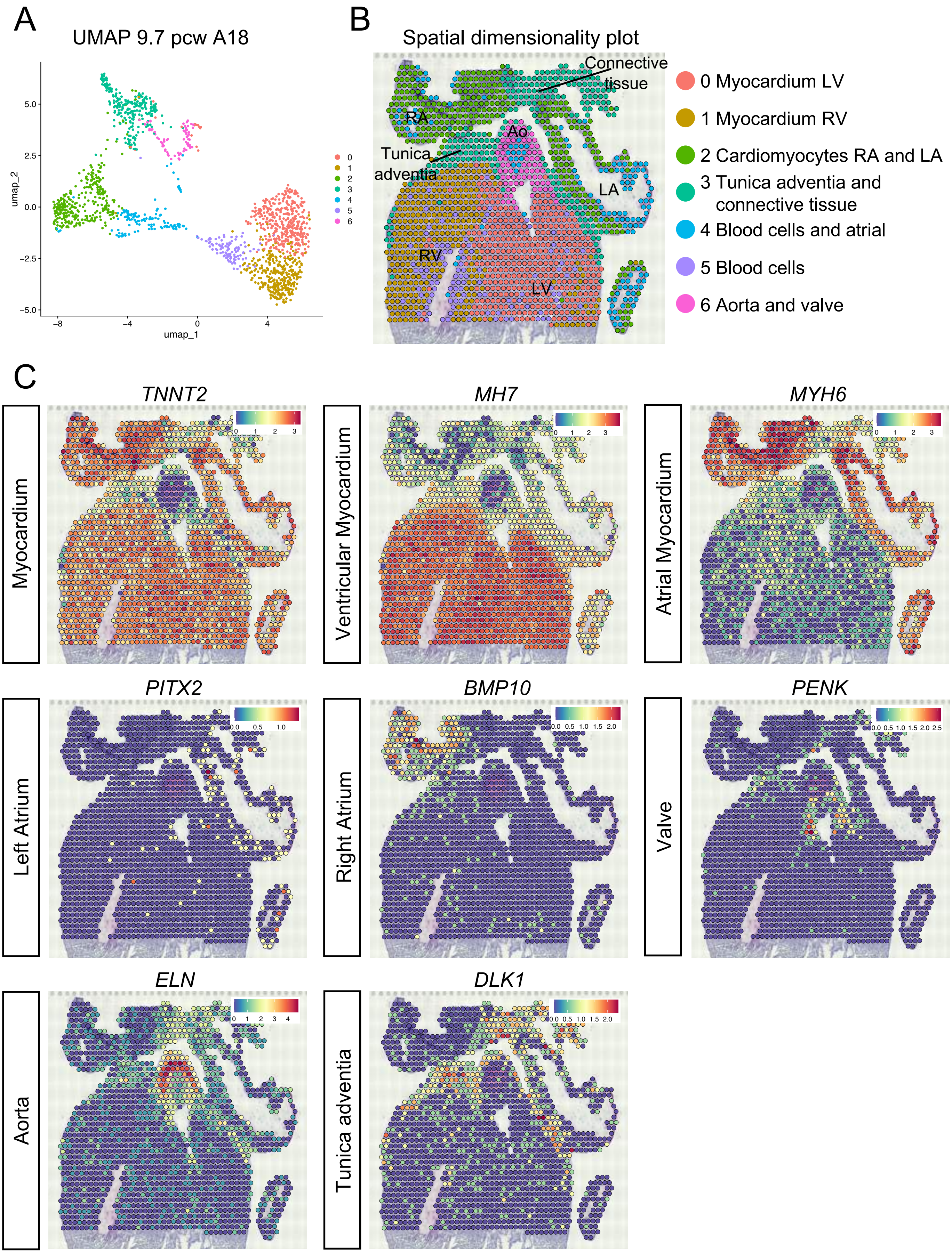
A
UMAP 9.7 pcw A18
umap_2
umap_1
B
Spatial dimensionality plot
Connective tissue
RA
Ao
Tunica adventia
LA
RV
LV
0 Myocardium LV
1 Myocardium RV
2 Cardiomyocytes RA and LA
3 Tunica adventia and connective tissue
4 Blood cells and atrial
5 Blood cells
6 Aorta and valve
C
Myocardium
TNNT2
Ventricular Myocardium
MH7
Atrial Myocardium
MYH6
Left Atrium
PITX2
Right Atrium
BMP10
Valve
PENK
Aorta
ELN
Tunica adventia
DLK1

Fig S7

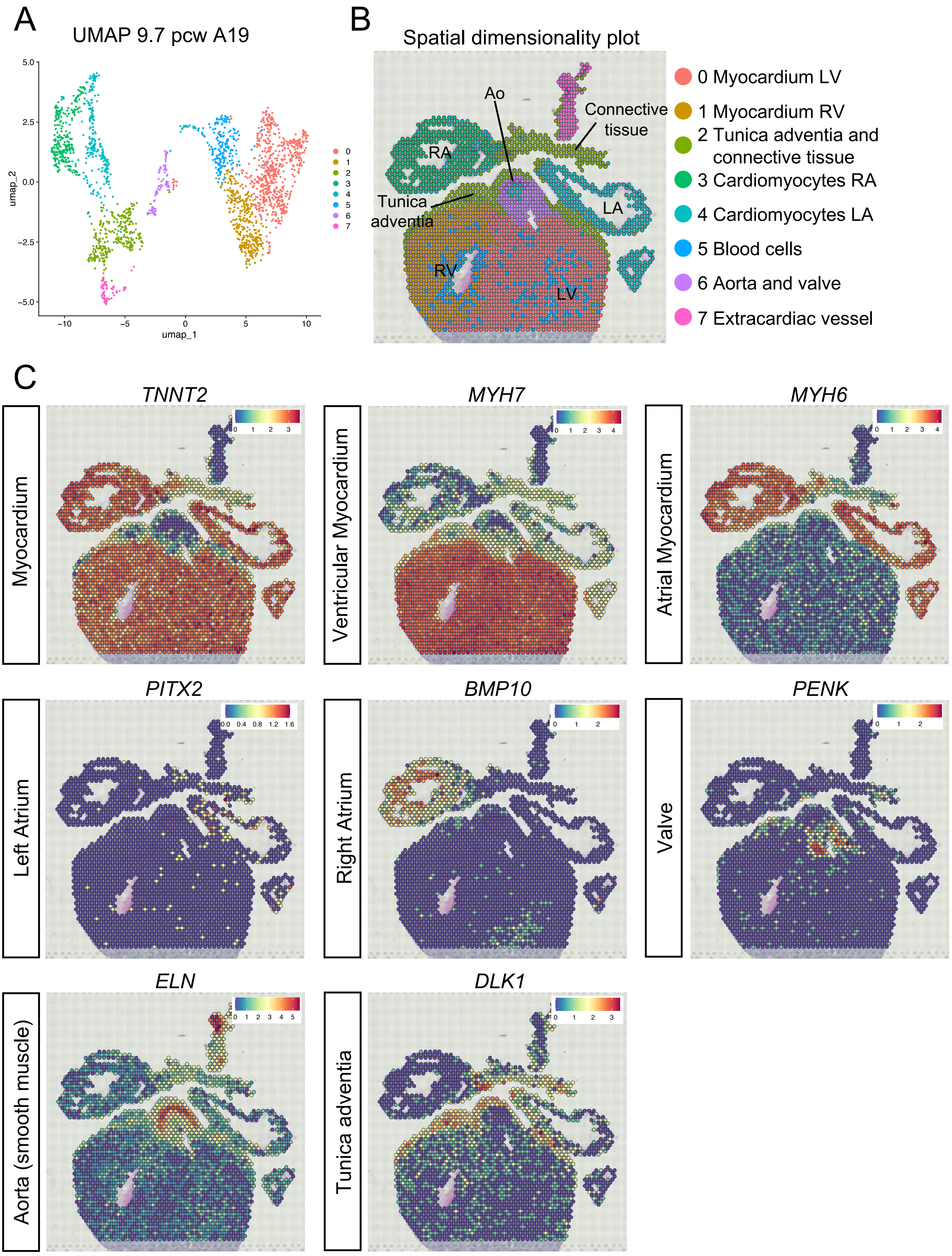

A
UMAP 9.7 pcw A19
umap_2
umap_1
0
1
2
3
4
5
6
7
B
Spatial dimensionality plot
Ao
Connective tissue
RA
LA
Tunica adventia
RV
LV
0 Myocardium LV
1 Myocardium RV
2 Tunica adventia and connective tissue
3 Cardiomyocytes RA
4 Cardiomyocytes LA
5 Blood cells
6 Aorta and valve
7 Extracardiac vessel
C
Myocardium
TNNT2
Ventricular Myocardium
MYH7
Atrial Myocardium
MYH6
Left Atrium
PITX2
Right Atrium
BMP10
Valve
PENK
Aorta (smooth muscle)
ELN
Tunica adventia
DLK1


Fig S8

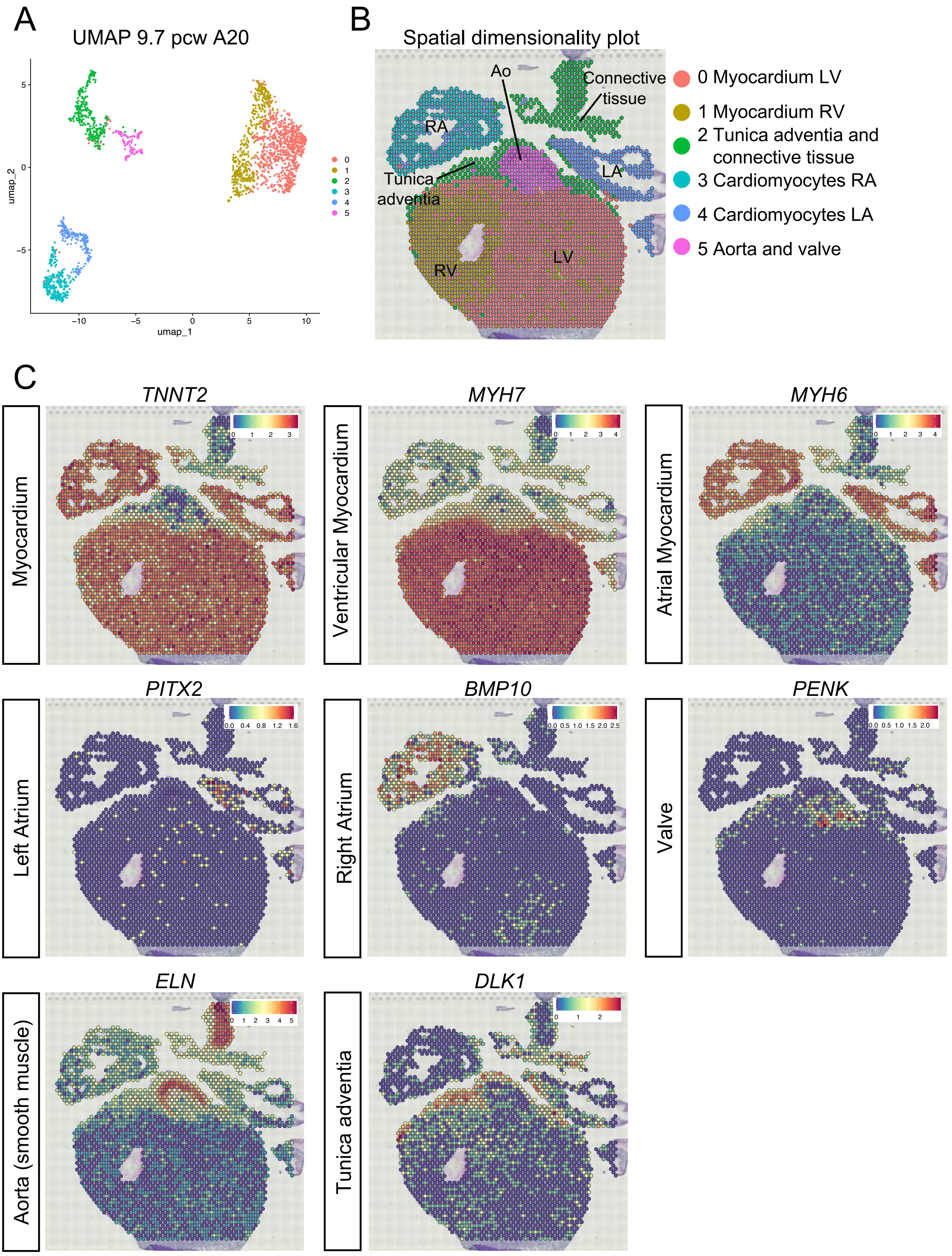
A
UMAP 9.7 pcw A20
umap_2
umap_1
B
Spatial dimensionality plot
Ao
Connective
tissue
RA
Tunica
adventia
LA
RV
LV
0 Myocardium LV
1 Myocardium RV
2 Tunica adventia and
connective tissue
3 Cardiomyocytes RA
4 Cardiomyocytes LA
5 Aorta and valve
C
TNNT2
MYH7
MYH6
Myocardium
Ventricular Myocardium
Atrial Myocardium
PITX2
BMP10
PENK
Left Atrium
Right Atrium
Valve
ELN
DLK1
Aorta (smooth muscle)
Tunica adventia

Fig S9

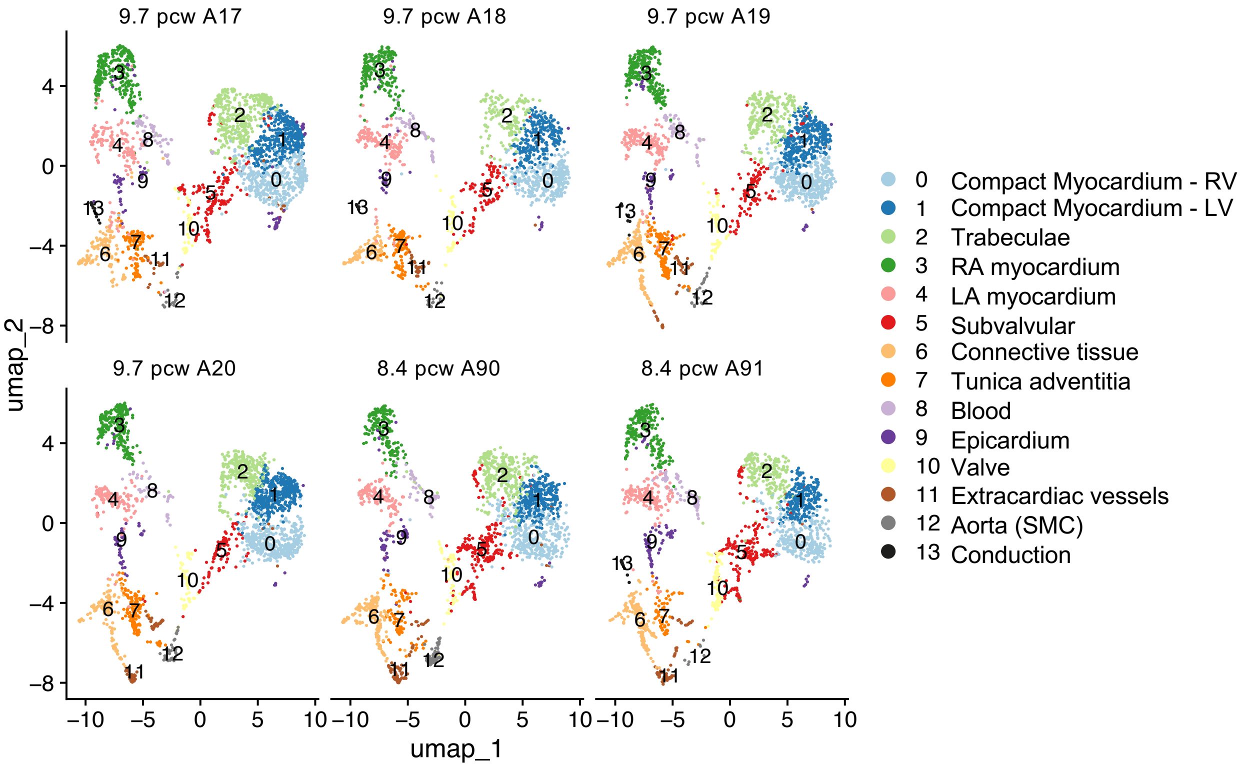
9.7 pcw A17
9.7 pcw A18
9.7 pcw A19
9.7 pcw A20
8.4 pcw A90
8.4 pcw A91
umap_2
umap_1
0 Compact Myocardium - RV
1 Compact Myocardium - LV
2 Trabeculae
3 RA myocardium
4 LA myocardium
5 Subvalvular
6 Connective tissue
7 Tunica adventitia
8 Blood
9 Epicardium
10 Valve
11 Extracardiac vessels
12 Aorta (SMC)
13 Conduction

Fig S10

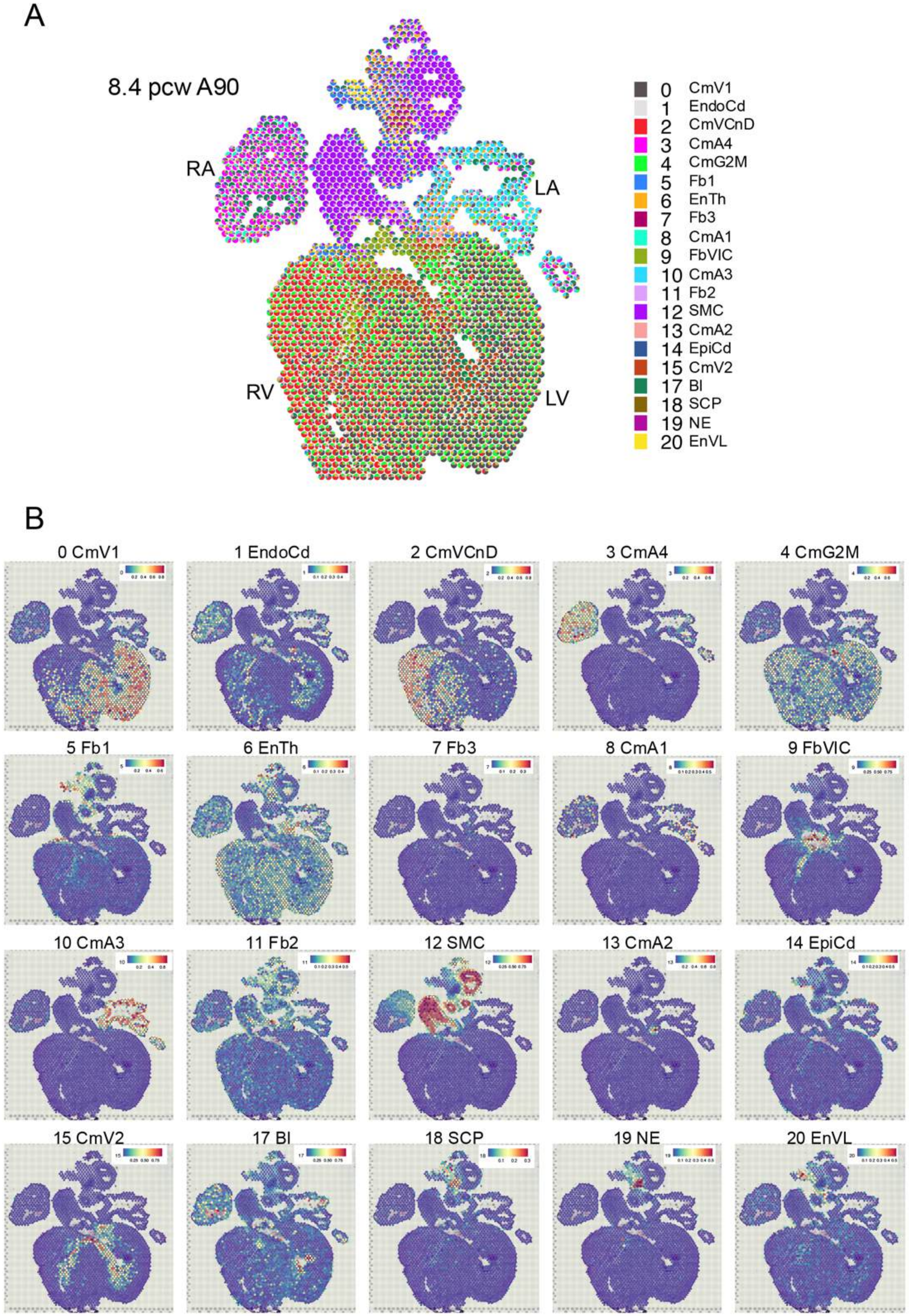
A
8.4 pcw A90
RA
LA
RV
LV
0 CmV1
1 EndoCd
2 CmVCnD
3 CmA4
4 CmG2M
5 Fb1
6 EnTh
7 Fb3
8 CmA1
9 FbVIC
10 CmA3
11 Fb2
12 SMC
13 CmA2
14 EpiCd
15 CmV2
17 Bl
18 SCP
19 NE
20 EnVL
B
0 CmV1
1 EndoCd
2 CmVCnD
3 CmA4
4 CmG2M
5 Fb1
6 EnTh
7 Fb3
8 CmA1
9 FbVIC
10 CmA3
11 Fb2
12 SMC
13 CmA2
14 EpiCd
15 CmV2
17 Bl
18 SCP
19 NE
20 EnVL

Fig S11

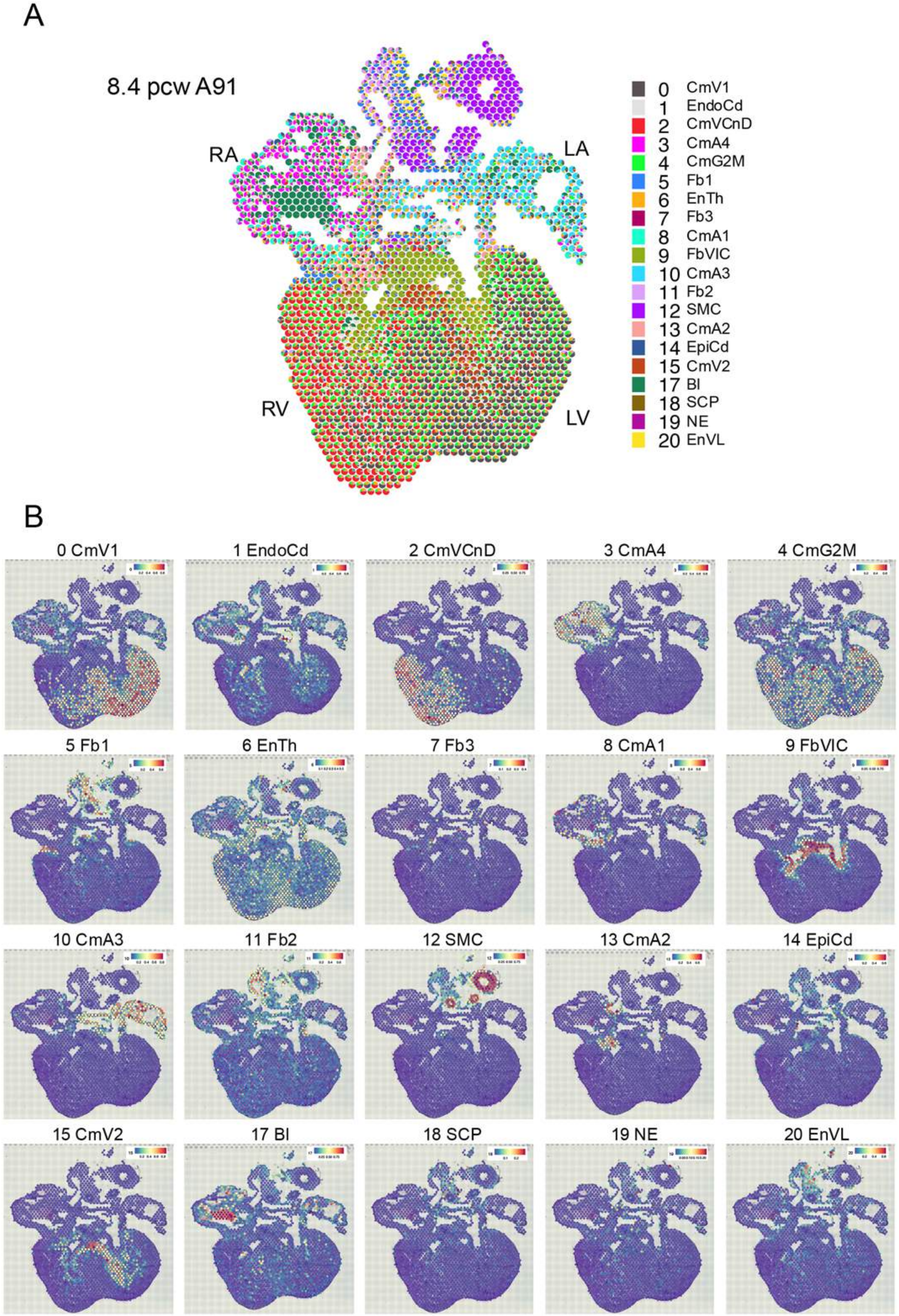
A
8.4 pcw A91
RA
LA
RV
LV
0 CmV1
1 EndoCd
2 CmVCnD
3 CmA4
4 CmG2M
5 Fb1
6 EnTh
7 Fb3
8 CmA1
9 FbVIC
10 CmA3
11 Fb2
12 SMC
13 CmA2
14 EpiCd
15 CmV2
17 Bl
18 SCP
19 NE
20 EnVL
B
0 CmV1
1 EndoCd
2 CmVCnD
3 CmA4
4 CmG2M
5 Fb1
6 EnTh
7 Fb3
8 CmA1
9 FbVIC
10 CmA3
11 Fb2
12 SMC
13 CmA2
14 EpiCd
15 CmV2
17 Bl
18 SCP
19 NE
20 EnVL

Fig S12

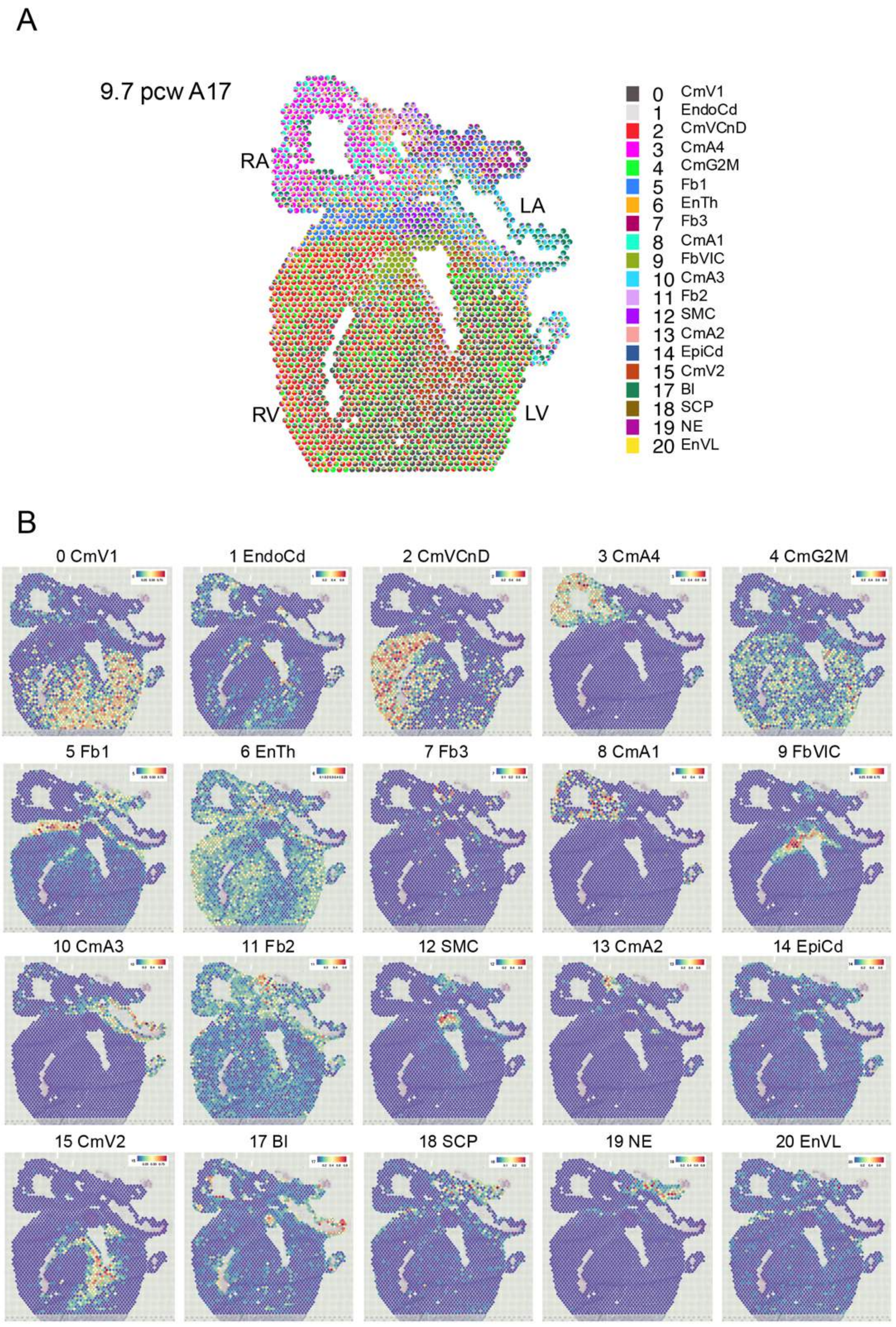
A
9.7 pcw A17
RA
LA
RV
LV
0 CmV1
1 EndoCd
2 CmVCnD
3 CmA4
4 CmG2M
5 Fb1
6 EnTh
7 Fb3
8 CmA1
9 FbVIC
10 CmA3
11 Fb2
12 SMC
13 CmA2
14 EpiCd
15 CmV2
17 Bl
18 SCP
19 NE
20 EnVL
B
0 CmV1
1 EndoCd
2 CmVCnD
3 CmA4
4 CmG2M
5 Fb1
6 EnTh
7 Fb3
8 CmA1
9 FbVIC
10 CmA3
11 Fb2
12 SMC
13 CmA2
14 EpiCd
15 CmV2
17 Bl
18 SCP
19 NE
20 EnVL

Fig S13

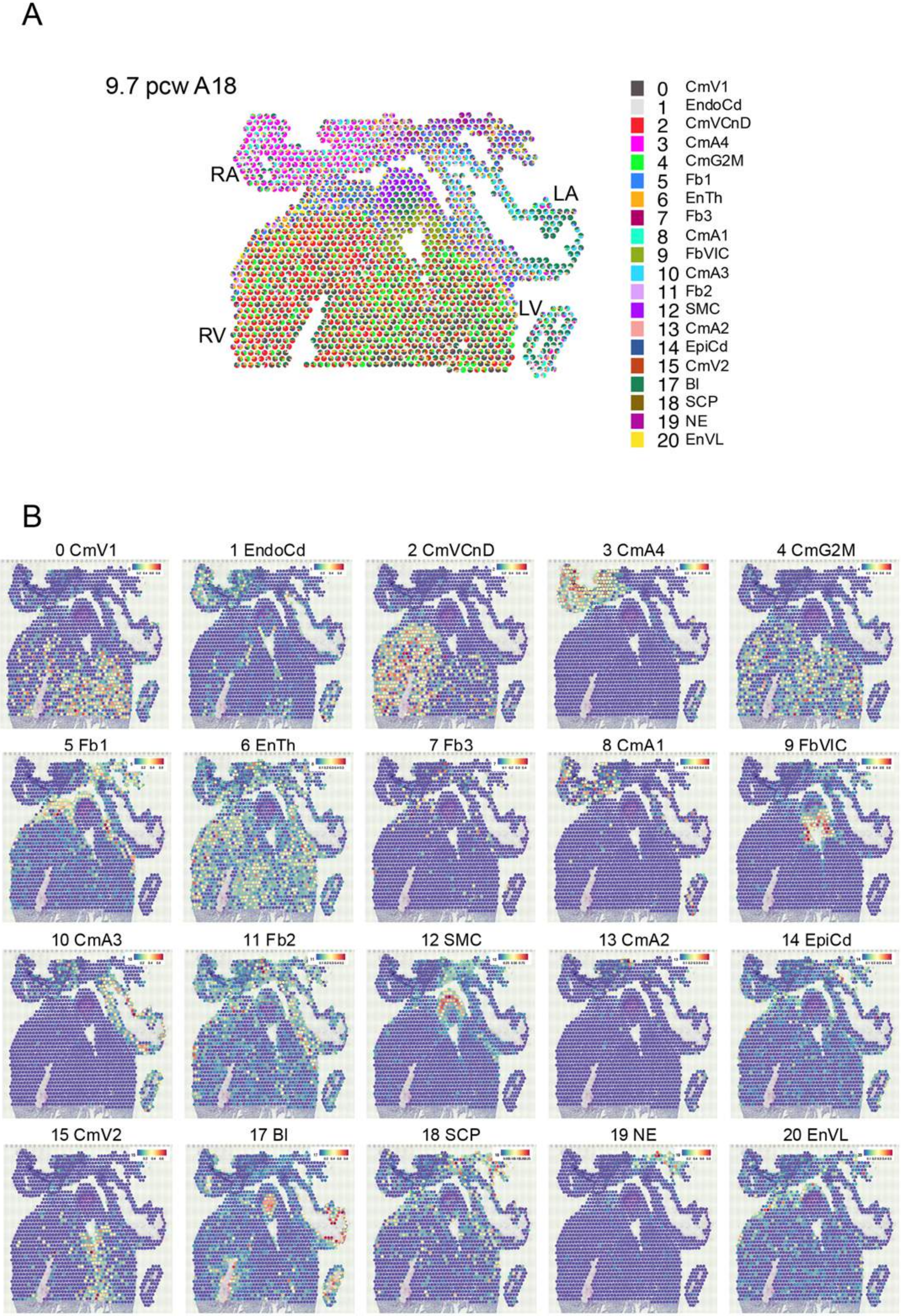
A
9.7 pcw A18
RA
LA
LV
RV
0 CmV1
1 EndoCd
2 CmVCnD
3 CmA4
4 CmG2M
5 Fb1
6 EnTh
7 Fb3
8 CmA1
9 FbVIC
10 CmA3
11 Fb2
12 SMC
13 CmA2
14 EpiCd
15 CmV2
17 Bl
18 SCP
19 NE
20 EnVL
B
0 CmV1
1 EndoCd
2 CmVCnD
3 CmA4
4 CmG2M
5 Fb1
6 EnTh
7 Fb3
8 CmA1
9 FbVIC
10 CmA3
11 Fb2
12 SMC
13 CmA2
14 EpiCd
15 CmV2
17 Bl
18 SCP
19 NE
20 EnVL

Fig S14

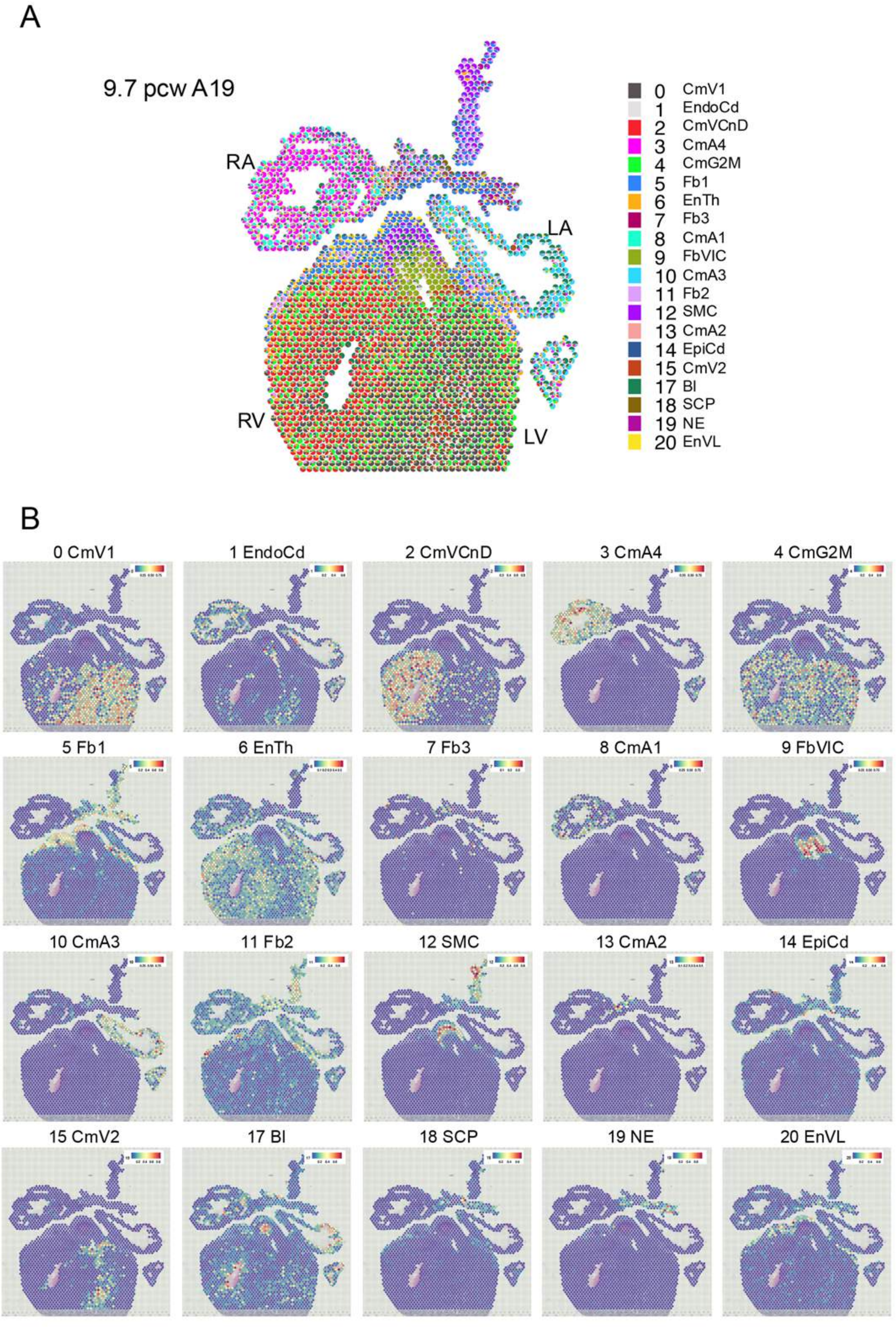
A
9.7 pcw A19
RA
LA
RV
LV
0 CmV1
1 EndoCd
2 CmVCnD
3 CmA4
4 CmG2M
5 Fb1
6 EnTh
7 Fb3
8 CmA1
9 FbVIC
10 CmA3
11 Fb2
12 SMC
13 CmA2
14 EpiCd
15 CmV2
17 Bl
18 SCP
19 NE
20 EnVL
B
0 CmV1
1 EndoCd
2 CmVCnD
3 CmA4
4 CmG2M
5 Fb1
6 EnTh
7 Fb3
8 CmA1
9 FbVIC
10 CmA3
11 Fb2
12 SMC
13 CmA2
14 EpiCd
15 CmV2
17 Bl
18 SCP
19 NE
20 EnVL

Fig S15

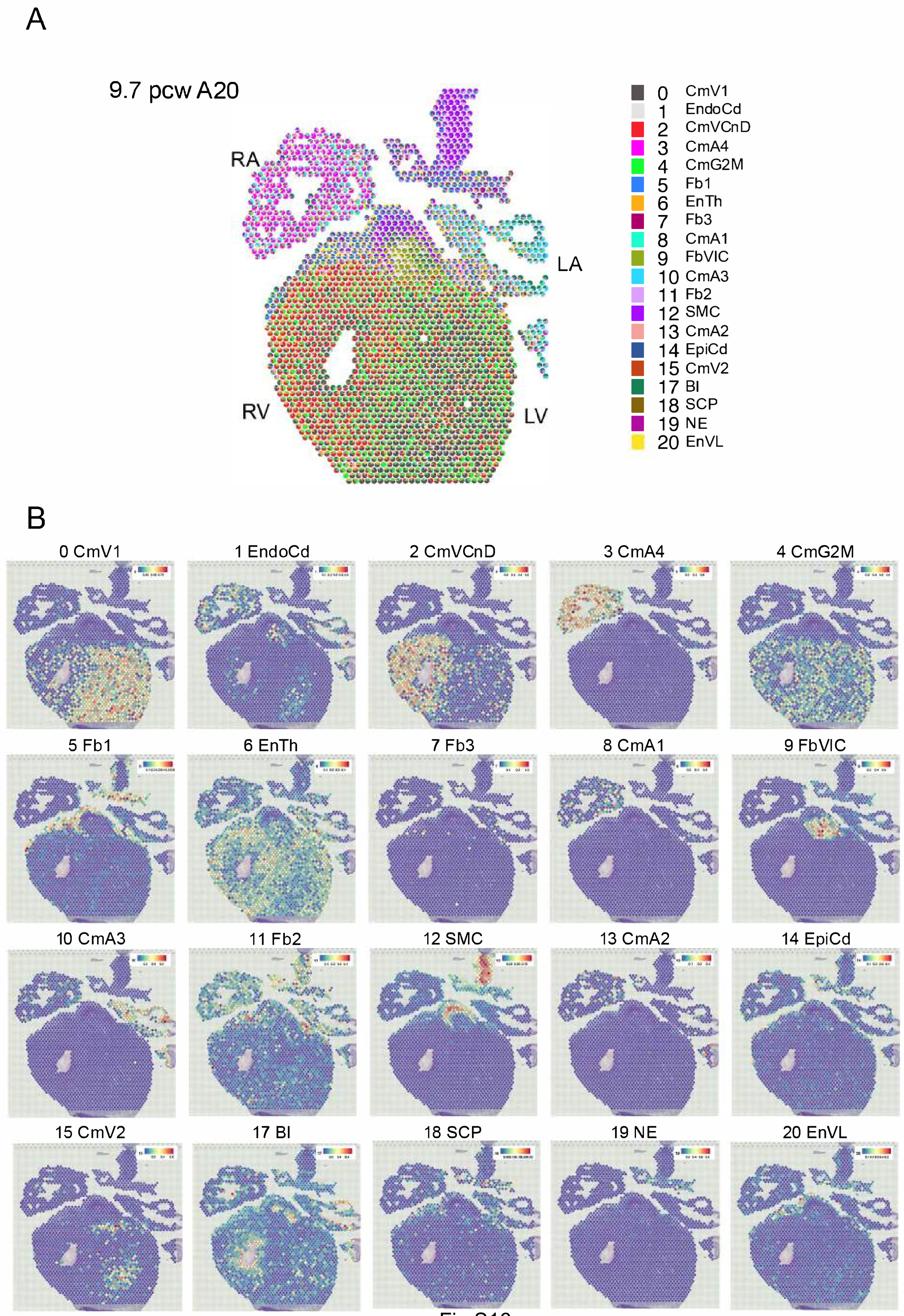
A
9.7 pcw A20
RA
LA
RV
LV
0 CmV1
1 EndoCd
2 CmVCnD
3 CmA4
4 CmG2M
5 Fb1
6 EnTh
7 Fb3
8 CmA1
9 FbVIC
10 CmA3
11 Fb2
12 SMC
13 CmA2
14 EpiCd
15 CmV2
17 BI
18 SCP
19 NE
20 EnVL
B
0 CmV1
1 EndoCd
2 CmVCnD
3 CmA4
4 CmG2M
5 Fb1
6 EnTh
7 Fb3
8 CmA1
9 FbVIC
10 CmA3
11 Fb2
12 SMC
13 CmA2
14 EpiCd
15 CmV2
17 BI
18 SCP
19 NE
20 EnVL

Fig S16

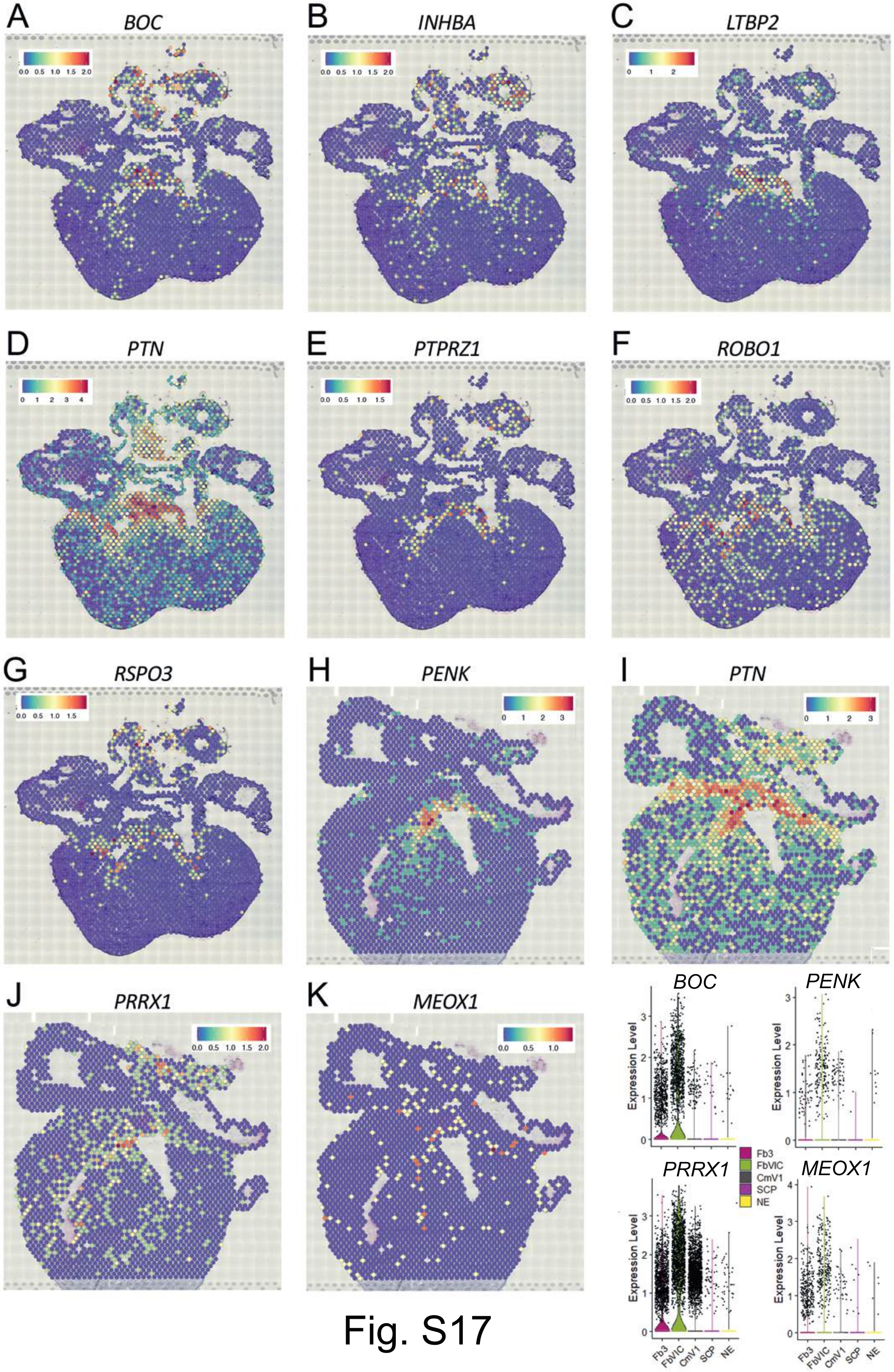

A
BOC
B
INHBA
C
LTBP2
D
PTN
E
PTPRZ1
F
ROBO1
G
RSPO3
H
PENK
I
PTN
J
PRRX1
K
MEOX1
BOC
PENK
PRRX1
MEOX1
Expression Level
Fb3
FbVIC
CmV1
SCP
NE


Fig. S17

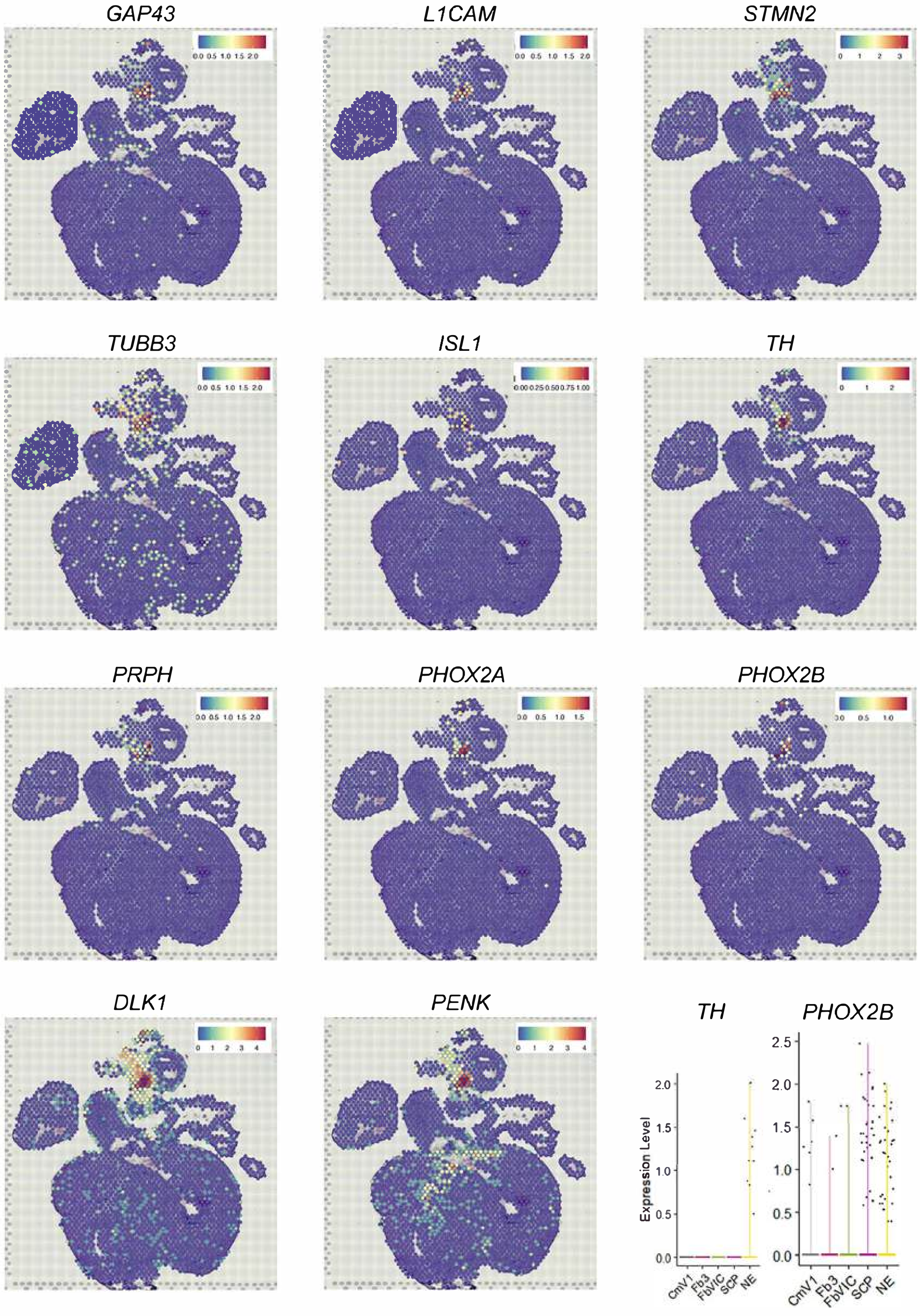
GAP43
L1CAM
STMN2
TUBB3
ISL1
TH
PRPH
PHOX2A
PHOX2B
DLK1
PENK
TH
PHOX2B
Expression Level
CmV1
Fb3
FbVIC
SCP
NE

Fig S18

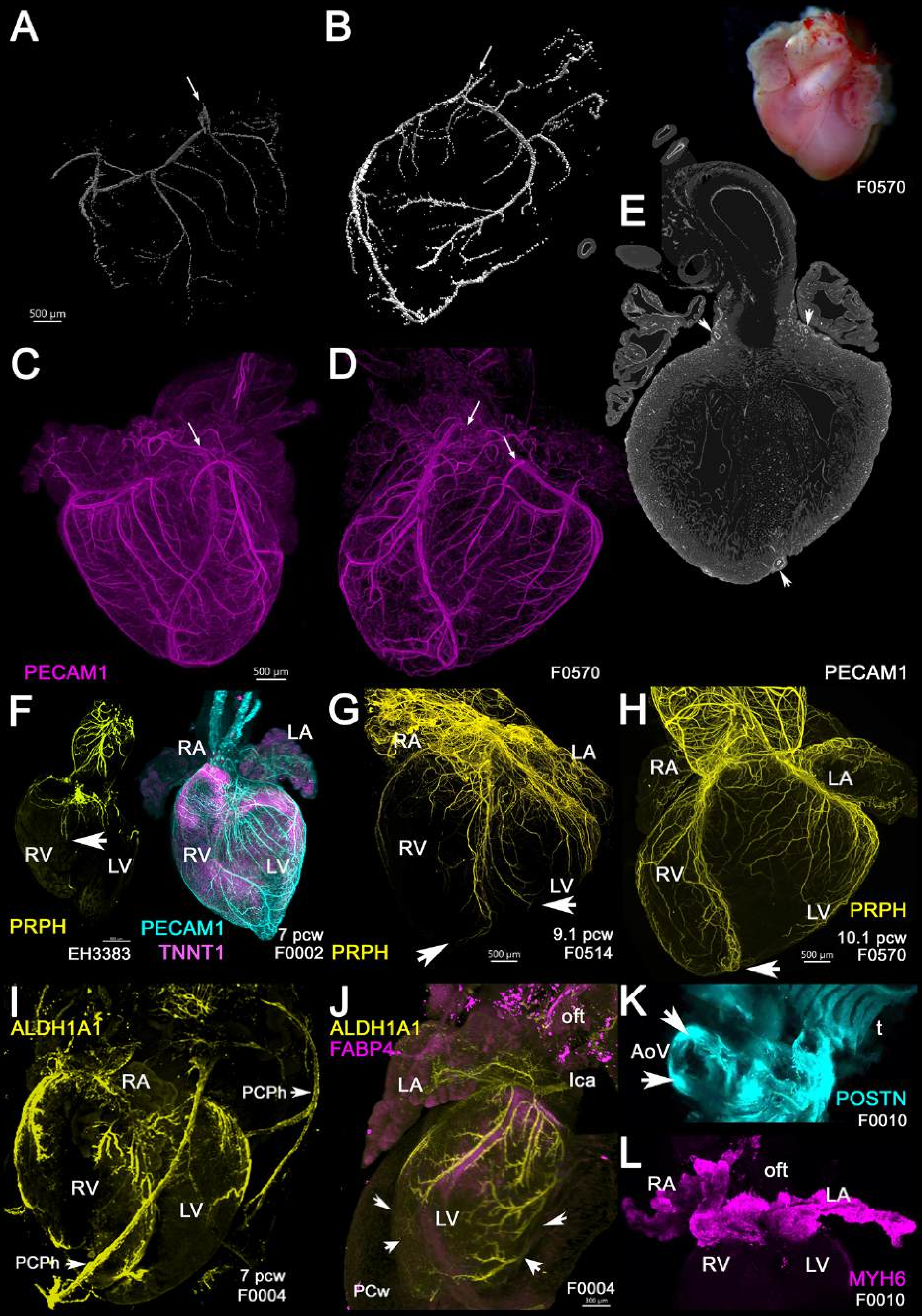
A
B
500 µm
F0570
E
C
D
PECAM1
500 µm
F0570
PECAM1
F
RA
LA
RV
LV
RV
LV
PRPH
EH3383
PECAM1
TNNT1
7 pcw
F0002
G
RA
LA
RV
LV
PRPH
500 µm
9.1 pcw
F0514
H
RA
LA
RV
LV
PRPH
10.1 pcw
500 µm
F0570
I
ALDH1A1
RA
PCPh
RV
LV
PCPh
7 pcw
F0004
J
ALDH1A1
FABP4
oft
LA
lca
LV
PCw
F0004
K
AoV
t
POSTN
F0010
L
RA
oft
LA
RV
LV
MYH6
F0010

Fig S19

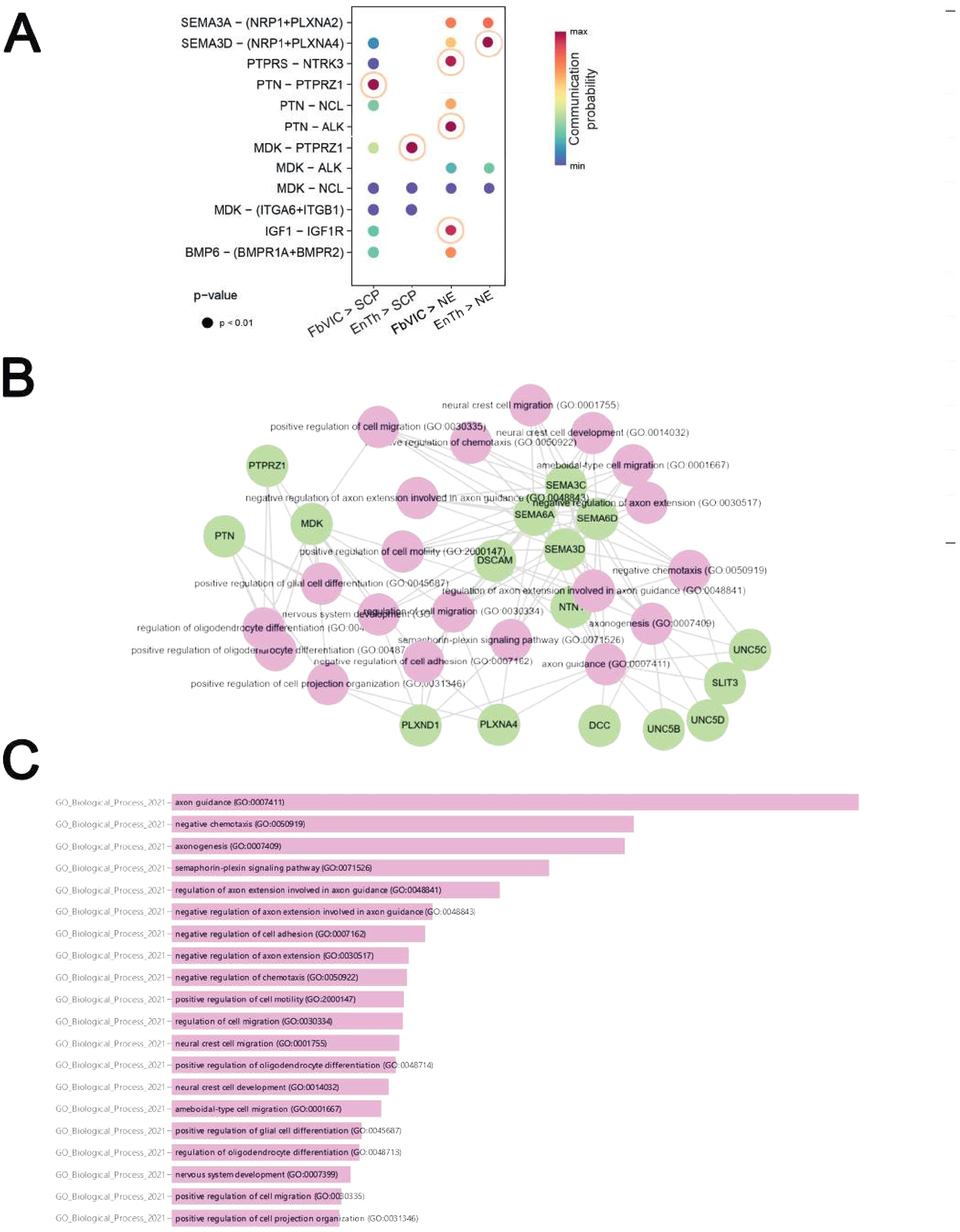
A
SEMA3A – (NRP1+PLXNA2)
SEMA3D – (NRP1+PLXNA4)
PTPRS – NTRK3
PTN – PTPRZ1
PTN – NCL
PTN – ALK
MDK – PTPRZ1
MDK – ALK
MDK – NCL
MDK – (ITGA6+ITGB1)
IGF1 – IGF1R
BMP6 – (BMPR1A+BMPR2)
FbVIC > SCP
EnTh > SCP
FbVIC > NE
EnTh > NE
p-value
p < 0.01
max
min
Communication probability
B
neural crest cell migration (GO:0001755)
positive regulation of cell migration (GO:0030335)
neural crest cell development (GO:0014032)
ameboidal-type cell migration (GO:0001667)
negative regulation of axon extension involved in axon guidance (GO:0048843)
negative regulation of axon extension (GO:0030517)
positive regulation of cell motility (GO:2000147)
negative chemotaxis (GO:0050919)
positive regulation of glial cell differentiation (GO:0045687)
regulation of axon extension involved in axon guidance (GO:0048841)
regulation of cell migration (GO:0030334)
axonogenesis (GO:0007409)
semaphorin-plexin signaling pathway (GO:0071526)
negative regulation of cell adhesion (GO:0007162)
axon guidance (GO:0007411)
positive regulation of cell projection organization (GO:0031346)
PTPRZ1
PTN
MDK
SEMA3C
SEMA6A
SEMA6D
SEMA3D
DSCAM
UNC5C
SLIT3
UNC5D
UNC5B
DCC
PLXNA4
PLXND1
C
GO_Biological_Process_2021
axon guidance (GO:0007411)
negative chemotaxis (GO:0050919)
axonogenesis (GO:0007409)
semaphorin-plexin signaling pathway (GO:0071526)
regulation of axon extension involved in axon guidance (GO:0048841)
negative regulation of axon extension involved in axon guidance (GO:0048843)
negative regulation of cell adhesion (GO:0007162)
negative regulation of axon extension (GO:0030517)
negative regulation of chemotaxis (GO:0050922)
positive regulation of cell motility (GO:2000147)
regulation of cell migration (GO:0030334)
neural crest cell migration (GO:0001755)
positive regulation of oligodendrocyte differentiation (GO:0048714)
neural crest cell development (GO:0014032)
ameboidal-type cell migration (GO:0001667)
positive regulation of glial cell differentiation (GO:0045687)
regulation of oligodendrocyte differentiation (GO:0048713)
nervous system development (GO:0007399)
positive regulation of cell migration (GO:0030335)
positive regulation of cell projection organization (GO:0031346)

Fig S20

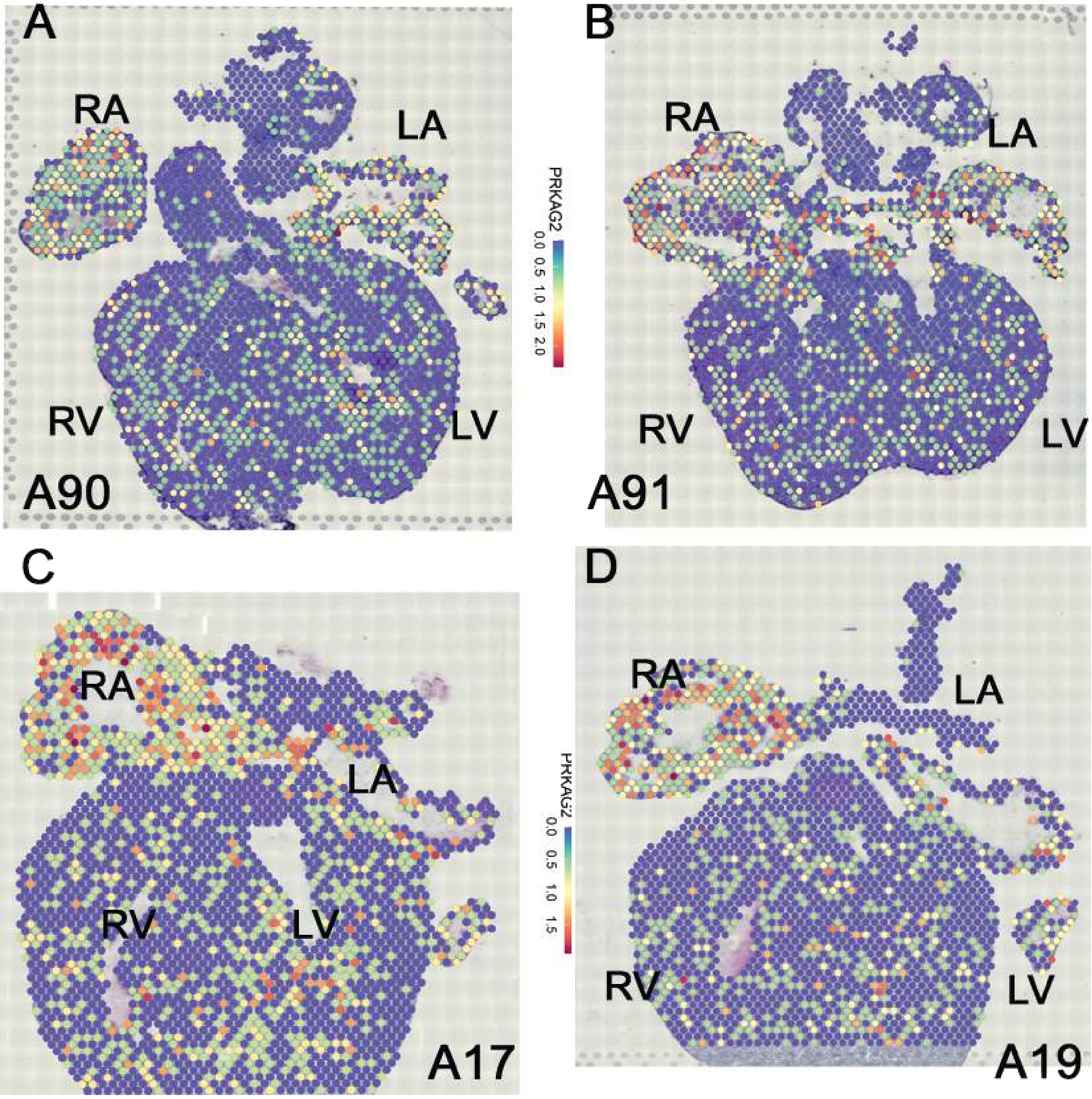
A
RA
LA
PRKAG2
0.0 0.5 1.0 1.5 2.0
RV
LV
A90
B
RA
LA
RV
LV
A91
C
RA
LA
RV
LV
A17
D
RA
LA
PRKAG2
0.0 0.5 1.0 1.5
RV
LV
A19

Fig S21
